# Cis-Attenuation of Pathogenic *Scn8a* Variant Causing Childhood Epilepsy Reveals Opposing Transcriptional Programs Driving Na_V_1.6 Gain and Loss of Function Phenotypes

**DOI:** 10.64898/2026.09.08.750173

**Authors:** Michael F. Hammer, Erfan Bahramnejad

**Author notes:** Corresponding author: Michael F Hammer, BIO5 Institute University of Arizona Tucson AZ 85721 USA.

## Abstract

Pathogenic variants in *SCN8A*, encoding the voltage-gated sodium channel Na_V_1.6, cause disease through opposing gain-of-function (GoF) and loss-of-function (LoF) mechanisms, yet the downstream cellular programs that distinguish these directions have not been resolved within a single genetic system. We used an isogenic *Scn8a* allelic series in which a *cis*-acting modifier produces a stepwise reduction of Na_V_1.6 on the N1768D background, isolating GoF, rescued, and LoF states on a shared background, and profiled the hippocampal transcriptome with pathway-level inference. The gain- and loss-of-function extremes engaged mechanistically opposite programs: an active fibro-inflammatory injury cascade in the seizing brain versus suppression of neuronal-signaling and growth programs, including cAMP Response Element-Binding signaling (CREB) in neurons, in the LoF state. The injury core was conserved across genetic backgrounds, appeared only after seizures rather than tracking channel dose, and was durably suppressed in the rescued genotype without erosion after seizure onset. Genetic rescue and the repurposed angiotensin-receptor blocker candesartan converged on the same injury program through distinct routes, and the same program was engaged in an α-synuclein model of Parkinson’s disease and in human epileptic tissue. These findings show that the two clinical directions of *SCN8A* disease have distinct tissue-level correlates and call for opposite therapeutic logic, channel or injury-cascade suppression for GoF and restoration of channel output for LoF, and identify a conserved, seizure-driven injury core as a tractable cross-disease target.

## 1. Introduction

The spectrum of phenotypes associated with monogenic channelopathies is striking. Pathogenic variants in genes encoding voltage-gated sodium channels (VGSCs) underlie muscle, cardiac, and neurological diseases, the presentations of which are often dictated by distinct gain-of-function (GoF) or loss-of-function (LoF) channel properties (1–3). However, the downstream cellular mechanisms generating these diverse phenotypes for a given VGSC remain poorly characterized. This gap in knowledge is due, in part, to a lack of preclinical models capable of isolating both GoF and LoF phenotypes on an otherwise identical, isogenic background. To address this, we utilized a novel *Scn8a* allelic series featuring a stepwise dose reduction of Na_V_1.6 expression, built upon the established *Scn8a*^N1768D^ GoF platform (4, 5).

GoF variants in *SCN8A*, encoding the VGSC Na_V_1.6, cause a severe and frequently drug-resistant developmental and epileptic encephalopathy (6–8). Because Na_V_1.6 is essential for normal neuronal firing (9), both excess (GoF) and deficiency (LoF) of channel activity are pathological — *SCN8A* behaves as a “Goldilocks” gene whose activity must be held within a narrow functional window (10). Emerging gene therapies that lower Na_V_1.6 must therefore titrate it precisely: too little correction leaves seizures, too much produces LoF deficits (11–13). Defining the protective level of Na_V_1.6, and the downstream molecular program it engages, is a central problem for *SCN8A*-related epilepsy. Clinically, the two directions diverge: GoF variants typically cause early-onset, seizure-dominated Developmental Epileptic Encephalopathy (DEE), whereas LoF variants more often cause intellectual disability, movement disorder, or absence epilepsy with a distinct comorbidity profile—divergent presentations whose brain-tissue mechanisms have not been compared within a single model (14–16).

A key unresolved question is whether the pathology of GoF *SCN8A* reflects the channel-level change itself or the downstream injury cascades that recurrent seizures set in motion (2, 17, 18). The distinction is therapeutically consequential: if seizures drive a self-propagating injury program, that program, rather than the channel, may offer safer, more tractable targets. Resolving channel-versus-cascade requires a system in which Na_V_1.6 dose can be varied and the resulting transcriptional state read out across disease, rescue, and pharmacologic intervention.

We recently identified a natural genetic modifier that provides such a system (companion paper, (19). A *de novo* 2-bp deletion arose in *cis* with the *Scn8a*-N1768D GoF allele and, through a frameshift and predicted nonsense-mediated decay (20, 21), functionally nullifies the mutant channel on its chromosome. In *Scn8a*-N1768D homozygotes, the number of modifier (L) copies titrates mutant Na_V_1.6 in graded fashion and yields three discrete phenotypes (**Figure 1**): short-lived mice with no modifier copies (SL; D+/D+; near-wild-type mutant Na_V_1.6, severe seizures, death ∼P21), long-lived mice with one copy (LL; DL/D+; ∼30% lower *Scn8a*, delayed onset ∼P49, survival ∼P65), and hindlimb-paralysis mice with two copies (HP; DL/DL; ∼70% lower *Scn8a*, LoF paralysis, euthanasia ∼P17) (companion paper, (19). This allelic series is, in effect, a built-in Na_V_1.6 dose-response spanning excess, optimum, and deficiency and provides a natural handle on both the therapeutic window and the mechanism that separates protection from harm. Because its extremes are excess (SL, GoF) and deficiency (HP, LoF) of Na_V_1.6 on a single genetic background, the series also provides a within-animal contrast of gain- and loss-of-function transcriptional states in the same tissue—a comparison not previously available.

**Figure 1.**
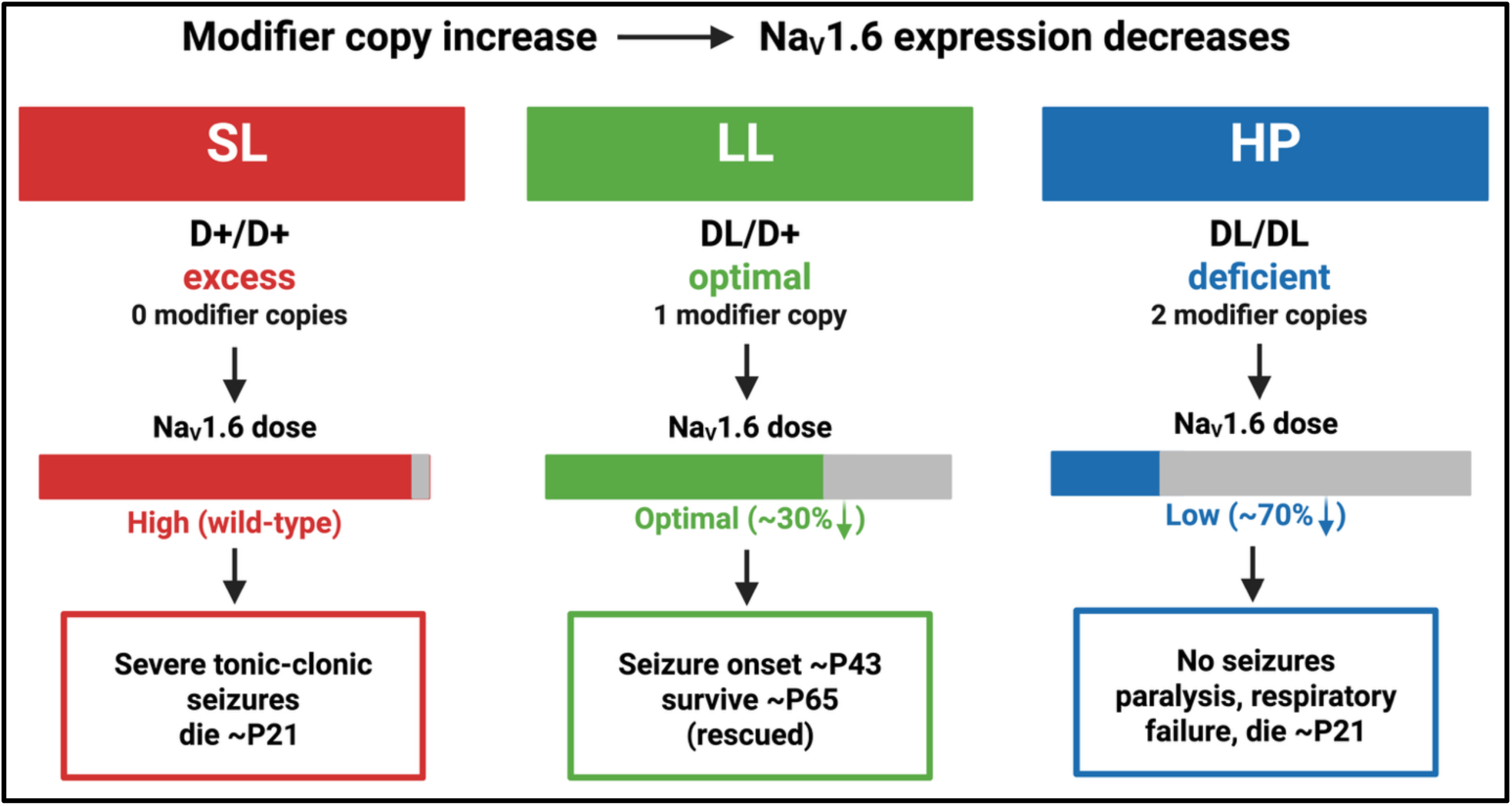
The *Scn8a*-N1768D modifier allelic series. A *de novo* 2-bp deletion in *cis* with *Scn8a*-N1768D (the modifier, L) titrates mutant Na_V_1.6 across three D/D genotypes: SL (D+/D+, no modifier copies; near-wild-type Na_V_1.6, severe seizures, death ∼P21), LL (DL/D+, one copy; ∼30% lower *Scn8a*, delayed onset, survival ∼P65), and HP (DL/DL, two copies; ∼70% lower *Scn8a*, LoF, euthanasia ∼P17). Both excess (SL) and deficiency (HP) are lethal; the intermediate dose (LL) is protective. Created in BioRender. Bahramnejad, E. (2026) https://BioRender.com/g7hxbhz.

Beyond the modifier series, the same model affords a broader comparative panel: GoF disease on two genetic backgrounds (C57BL/6J [B6] and the congenic C3H/HeJ·C57BL/6J [C3H·B6]), heterozygous and homozygous channel dose, pre- and post-seizure states, as well as a parallel cohort that demonstrated pharmacologic rescue with the angiotensin-receptor blocker candesartan (17). Read together, these contrasts can separate the stereotyped, seizure-associated injury program from background- and dose-specific effects and can test whether genetic dose-reduction and pharmacologic intervention converge on shared downstream targets. To interpret them on a common footing, we apply a pathway-effect inference framework that annotates each enriched canonical pathway for its likely beneficial or detrimental contribution— an approach we have applied to candesartan-treated mice and to human temporal-lobe-epilepsy resections (17, 22), and one that is portable across datasets and, in principle, across neurological disorders that share injury-cascade biology.

Here we use this comparative system to define how the modifier reshapes the hippocampal transcriptome at the level of canonical pathways, and to place that function in the context of treated, untreated, and cross-background mice. We show that modifier dose produces a graded, global quieting of the transcriptome; that the two pathological extremes engage opposite hippocampal programs: an inflammatory and tissue-remodeling signature at the GoF extreme (SL) and a suppressed growth-factor and neuronal-signaling signature at the LoF extreme (HP); that a conserved, background-independent GoF injury core is activated wherever disease is expressed and specifically inverted by the modifier; that the protective state is durable across seizure onset; and that the modifier and candesartan converge on the same injury core by distinct transcriptomic strategies. Together these findings argue that the modifier acts upstream of a seizure-associated injury cascade, and they establish a comparative pathway-effect framework for inferring mechanism and nominating safer targets in *SCN8A*-related epilepsy and related disorders. By contrasting GoF and LoF states in one model, the series further offers a tissue-level correlate of the divergent clinical presentations of human *SCN8A* variants.

## 2. Materials and Methods

### 2.1. Animals and genotyping

Mouse lines, genotyping and phenotyping procedures are describing in Bahramenjad et al. (companion manuscript). The genotypes and phenotypes of the mice studied here are listed in **Table 1**. Briefly, The *Scn8a*-N1768D knock-in mutation was constructed on the C57BL/J background () and a congenic C3H.B6-*Scn8a*^N1768D^ (C3H.B6) line was created by by backcrossing a male heterozygote (D/+) in B6 colony to a C3H female wild-type (WT) mouse. In the F1 generation, D/+ siblings were mated to generate homozygous (D/D) mice in the F2 generation. These mice were then monitored for seizure onset, lifespan and seizure frequency. In the N6 (98.4%) line, we observed that homozygous D/D mice displayed three distinct phenotypes. D/D mice of all three phenotypes were monitored 24/7 using a video camera system to detect their first tonic-clonic seizure (TCs), record the number of TC events during their lifespan, and determine the time of their death (18). When animals were euthanized, euthanasia was performed by cervical dislocation without anesthesia. No anesthetics were administered in this study. All animal work was performed in the animal facility in the Thomas Keating Building at the University of Arizona, Tucson, AZ, USA, and was approved by the University of Arizona Institutional Animal Care and Use Committee (IACUC protocol #16-160).

**Table 1.** Genotype, background, age, seizure state, treatment, and pathway-analysis status of the thirteen hippocampal RNA-seq contrasts.

|  | IPA label | Geno-<br>type | Modifier<br>copies | Back-<br>ground | Age | Seizure<br>state | Treat-<br>ment | # Sig<br>pathways | \|z\|><br>=2 | z-<br>availability |
| --- | --- | --- | --- | --- | --- | --- | --- | --- | --- | --- |
| 1 | B6-HOM<br>post | D/D | 0 | B6 | ~P21 | post-TC | none | 488 | 127 | rich |
| 2 | B6-HET<br>post | D/+ | — | B6 | ~P100 | post-TC | none | 395 | 151 | rich |
| 3 | N7B-HET<br>pre | D/+ | — | C3H.B6<br>(N7B) | ~P45 | pre-TC | none | 18 | 0 | enrichment-<br>only |
| 4 | N7B-HET<br>post | D/+ | — | C3H.B6<br>(N7B) | ~P100 | post-TC | none | 134 | 18 | partial |
| 5 | SL-HOM | D/D | 0 | C3H.B6<br>(SLxHP) | ~P21 | post-TC | none | 512 | 82 | rich |
| 6 | LL-HOM | D/D | 1 | C3H.B6<br>(SLxHP) | ~P21 | pre-TC | none | 313 | 112 | rich |
| 7 | HP-HOM | D/D | 2 | C3H.B6<br>(SLxHP) | ~P21 | non-seizing | none | 228 | 61 | rich |
| 8 | 98.5 pre | D/D | 1 (LL) | N6 | ~P42 | pre-TC | none | 62 | 2 | enrichment-<br>only |
| 9 | 98.5 post | D/D | 1 (LL) | N6 | ~P65 | post-TC | none | 36 | 0 | enrichment-<br>only |
| 10 | B6-HOM-trt | D/D | 0 | B6 | P15–<br>20 | treated | CND 5 d | 28 | 0 | enrichment-<br>only |
| 11 | B6-HET<br>pre trt | D/+ | — | B6 | pre-TC | treated | CND | 84 | 1 | enrichment-<br>only |
| 12 | B6-Het pos<br>trt | D/+ | — | B6 | post-TC | treated | CND | 136 | 2 | enrichment-<br>only |
| 13 | B6 HET<br>LTS trt | D/+ | — | B6 | adult | treated | CND<br>long-term | 100 | 0 | enrichment-<br>only |

B6= C57L6/J; N7B= 7<sup>th</sup> generation backcross; N6= 6<sup>th</sup> generation backcross; CND= candesartan; P= postnatal day; HET= heterozygote; HOM+ homozygote; TC= Tonic-clonic seizures; pre= pre-TC; post= post-TC; trt= Treated.

### 2.2. Gene expression and pathway enrichment analysis

Hippocampal RNA-seq differential expression (each genotype versus age-matched wild-type [WT] littermates) (for details, see Bahramnejad et al. companion paper) was analyzed with Ingenuity Pathway Analysis (IPA; Qiagen), which returned, for each canonical pathway and contrast, an enrichment significance (−log_10_ p) and, where estimable, an activation z-score (**Table 1**). Thirteen contrasts spanned the *Scn8a*-N1768D homozygous (D/D) and heterozygous (D/+) genotypes across two genetic backgrounds (B6 and the congenic C3H·B6), two developmental windows (juvenile ∼P21; adult ∼P100), pre- and post-seizure states, and candesartan treatment (**Table 1**). The modifier is a *de novo* 2-bp deletion in *cis* with *Scn8a*-N1768D that nullifies the mutant allele on its chromosome (companion paper) (19); the primary comparison therefore used three *Scn8a*-N1768D homozygous (D/D) genotypes carrying graded mutant Na_V_1.6 dose in juvenile mice: SL (D+/D+, no modifier copies) at P21, LL (DL/D+, one copy) at P21-P25, and HP (DL/DL, two copies) at P18. Pathways were considered significant at p < 0.05 (−log_10_ p > 1.301); directional analyses additionally required |z| ≥ 2.0. For a given genotype contrast, a pathway was counted as directional only if it met both criteria in that contrast (significant enrichment and |z| ≥ 2.0); pathways with significant enrichment but no estimable activation z-score were retained as significant but non-directional. Activated and deactivated directional pathways were defined as directional pathways with |z| ≥ +2.0 and |z| ≤ −2.0, respectively; the net activation index was the number activated minus the number deactivated within the stated pathway set.

Activation z-scores were richly estimated for the untreated genotype contrasts (directional pathway counts in the full IPA output ranged from 82 to 151 per contrast; **Table 1**) but were largely unavailable for the candesartan-treated contrasts and for the long-lived N6 longitudinal pair (19); analyses involving the latter were therefore restricted to enrichment presence, as specified below.

### 2.3. Pathway-effect annotation and consistency check

Each pathway was annotated for its involvement in fifteen recurrent injury/repair “effects” — excitotoxicity (ETX), oxidative stress (OXS), neuroinflammation (INF), glial activation (GLA), blood–brain barrier integrity (BBB), fibrosis/scar formation (FBS), extracellular-matrix remodeling (ECM), secondary injury cascade (SIC), neurotransmission (NTM), synaptic plasticity (SYN), axon guidance (AXG), apoptosis/clearance (APO), neurogenesis/repair (NER), angiogenesis (ANG), and neurodegeneration (NDG) (**Table 2**, **Table S1**). This set reconciles the twelve- and thirteen- effect schemes used in our prior candesartan (CND) (17) and temporal-lobe-epilepsy (TLE) (22) studies. For each engaged effect we recorded an activation-valence (detrimental, beneficial, or context-dependent), assigned from current literature and made blind to the genotype z-scores to prevent confirmation bias. To verify reproducibility of this re-derivation, annotations were compared cell-by-cell against the published per-pathway effect matrices (Table S3 in (22); Table S1 in (22) for overlapping pathways; 98.6% of co-annotated effect calls were non-contradictory, the small residual reflecting a deliberately more context-explicit encoding. Pathways whose primary identity was a peripheral neoplasm were excluded, yielding 180 pathways significant and directional in at least one of SL/LL/HP (**Table S1**). Because many canonical-pathway and disease-function labels derive from oncology, where these programs were first characterized at the pathway level, retained pathways were interpreted at the level of their constituent genes and conserved cellular processes (e.g., inflammation, extracellular-matrix and fibrotic remodeling, angiogenesis, and growth-factor and cytoskeletal signaling) rather than by their organ- or tumor-specific names, and pathway direction was read in the context of CNS injury rather than neoplasia.

**Table 2.**
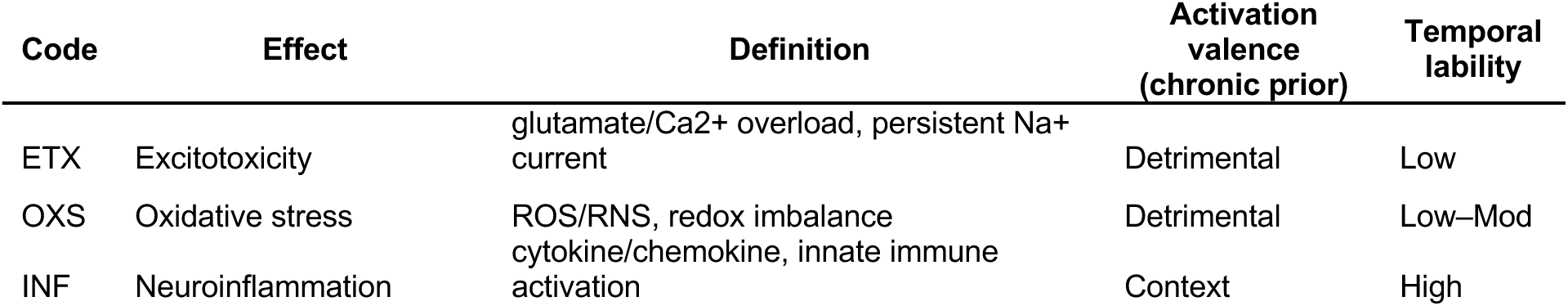
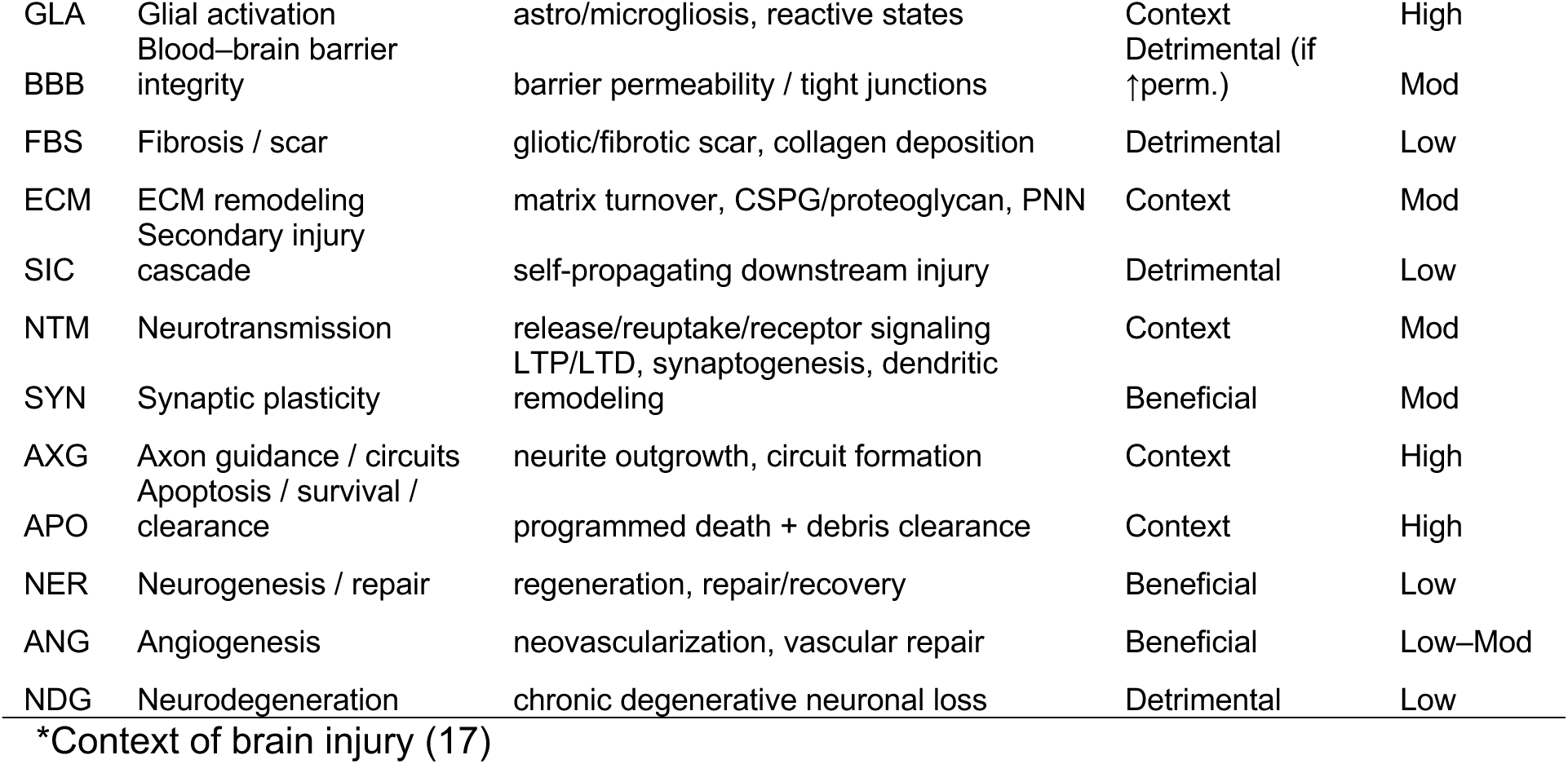
Fifteen recurrent injury and repair effects used to annotate canonical pathways, with definitions, chronic-state activation valence, and temporal lability*.

### 2.4. Genotype-specific realized scoring

Because the modifier genotypes occupy distinct disease states in the juvenile mice (i.e., SL post-seizure (chronic injury), LL pre-seizure (tonic-clonic onset ∼P49), and HP non-seizing with LoF paralysis), realized beneficial/detrimental tallies were computed under genotype-specific priors rather than a single chronic-stage assumption (**Table 3**). The chronic prior was applied to SL. For LL, growth and repair effects were treated as context (no active-injury demand) while injury- cascade effects retained their valence. For HP, neuronal-function effects (neurotransmission, synaptic plasticity, axon guidance) were scored such that their deactivation registered as detrimental (the LoF phenotype), while injury-cascade and growth effects were treated as context. A pathway’s net score was the number of detrimental minus beneficial effect-instances, with context calls tabulated separately.

**Table 3.** Genotype-specific biological states and scoring priors applied to realized beneficial and detrimental tallies.

| Genotype | Biological state at P21 | Scoring prior |
| --- | --- | --- |
| SL (D/D, 0 modifier copies) | post-seizure, chronic injury (~30 TCs) | Chronic: effects keep stored activation-valence |
| LL (D/D, 1 copy) | pre-seizure (onset ~P45), rescued | Developmental: growth/repair effects (SYN,NER,ANG,AXG,APO) → context; injury cascade keeps valence |
| HP (D/D, 2 copies) | non-seizure, loss-of-function paralysis | LOF: neuronal-function effects (NTM,SYN,AXG) → beneficial-when-activated (deactivation = detrimental); ETX stays detrimental; injury cascade + growth → context |

### 2.5. Cross-contrast analyses

Background-independence and apparent dose effects were assessed by comparing the set of activated injury pathways among the four z-rich untreated genotypes (B6 and C3H·B6 D/D; B6 and C3H·B6 D/+), with set overlap quantified by the Jaccard index (23). The durability of cascade suppression after seizure onset was assessed within a single long-lived N6 line sampled pre-TC (P42) and post-TC (P65); because these contrasts lacked activation z-scores, this comparison used enrichment presence only. Convergence of genetic and pharmacologic rescue was assessed for the conserved injury core by tabulating, in matched genotypes, how many core pathways lost enrichment after candesartan (loss of a significant difference from WT, i.e. normalization) versus how many were inverted (significantly deactivated) in the long-lived modifier genotype; the overlap defined a shared target program.

### 2.6. Network-level mechanistic inference

To complement the effect-level annotation, each phenotype-versus-WT contrast was additionally summarized at the network level using the graphical-summary (network-overview) function of Ingenuity Pathway Analysis (IPA), which integrates the most significant canonical pathways, predicted upstream regulators, and downstream functions for a contrast into a single directed network. SL was summarized at both FDR < 0.05 and FDR < 0.001 (to assess robustness of the dominant signature to threshold); LL and HP were summarized at FDR < 0.05. These overviews were used to characterize the dominant mechanistic signature of each genotype and were curated as in §1.2, with oncology-derived pathway and function labels interpreted at the level of constituent genes and conserved processes. Two artificial-intelligence aids were used in interpretation: the IPA network-summary function and a large-language-model assistant (Claude, Anthropic) for literature-based effect annotation. Outputs were treated as hypothesis-generating: all inferences were verified against primary literature and against the quantitative enrichment and activation-z results, which constitute the confirmatory backbone of the analysis. The authors take responsibility for all interpretations. For details on the differentially expressed genes (DEGs) inferred from the RNAseq data and that underlie the pathway enrichment analyses, see (17–19).

## 3. Results

### 3.1. Dose-dependent transcriptional quieting across the modifier series

Across the three modifier genotypes, 180 canonical pathways were significantly enriched and directional in at least one genotype. The most striking feature was independent of any effect annotation: the *direction* of pathway regulation tracked modifier dose. SL hippocampus was dominated by pathway activation (65 of 74 directional pathways activated; net activation index +56), whereas LL and HP were dominated by deactivation (4 of 104 and 1 of 57 activated; net indices −96 and −55) (**Figure 2**). Activation was almost entirely SL-specific (no activated pathway was shared with LL or HP), whereas deactivation was largely shared between LL and HP (**Figure 2C**). Thus, increasing modifier copy number and the corresponding graded reduction in *Scn8a*/Na_V_1.6 expression (SL > LL > HP) produced progressive, global quieting of the hippocampal transcriptome, most pronounced in LL.

**Figure 2.**
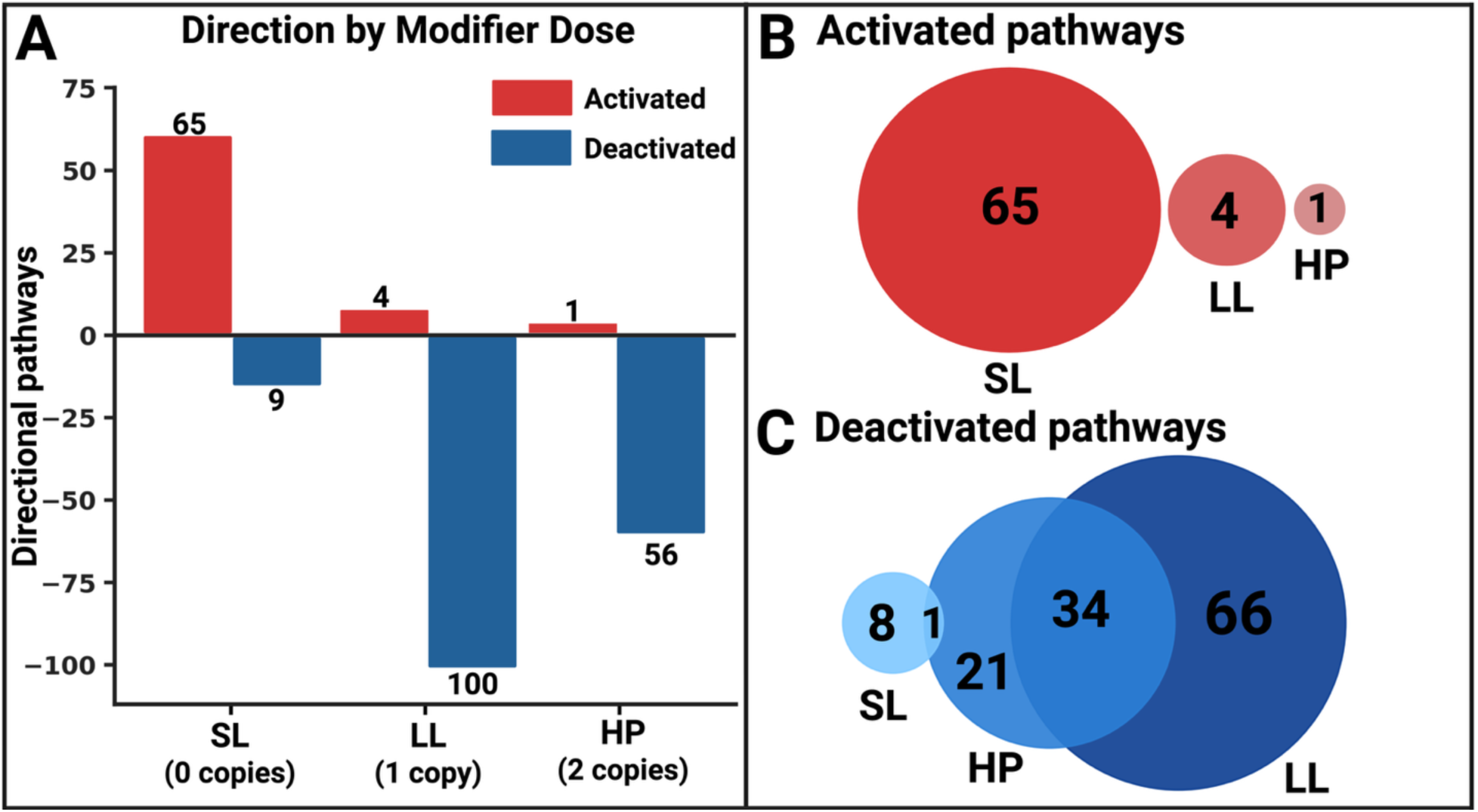
Dose-dependent direction asymmetry. (*A*) Activated and deactivated directional pathways (|z| ≥ 2) per modifier genotype; activation predominates in SL, deactivation in LL and HP. (*B*, *C*) Three-way overlap of (*B*) activated and (*C*) deactivated significant pathways across SL, LL, and HP. Activation is almost entirely SL-private; deactivation is largely shared between LL and HP. Created in BioRender. Bahramnejad, E. (2026) https://BioRender.com/5foyhtb.

Translating pathway direction into an inferred beneficial/detrimental balance required accounting for genotype state. Under a single chronic-injury assumption applied uniformly, LL paradoxically scored as the most detrimental genotype — an artifact arising because its deactivation of growth and repair programs was scored as lost reparative capacity despite the absence of active injury.

Under genotype-specific priors the balance resolved into three qualitatively distinct states: SL was net-neutral (a florid but balanced response), LL net-beneficial (suppression of injury tone without a functional penalty), and HP net-detrimental (deactivation of neuronal-function programs, consistent with its LoF phenotype) (**Figure 3**). The direction asymmetry is annotation-independent and robust; the beneficial/detrimental balance is an interpretive layer contingent on the priors and is treated accordingly.

**Figure 3.**
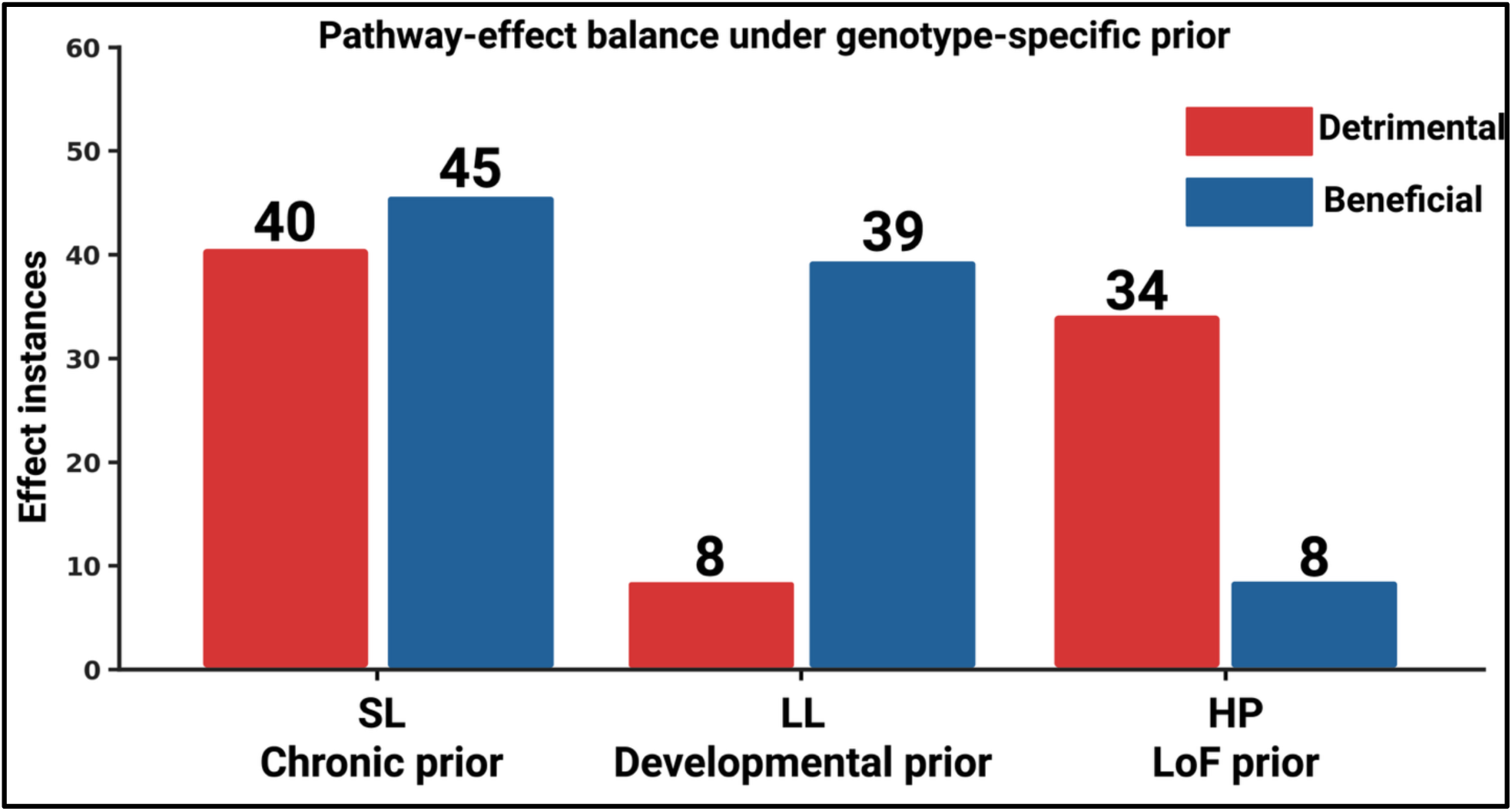
Pathway-effect balance under genotype-specific priors. Detrimental and beneficial effect-instances per genotype (context, dual-valence instances noted below the axis). The net detrimental–beneficial balance is most beneficial in LL and most detrimental in HP, with SL near-neutral. Created in BioRender. Bahramnejad, E. (2026) https://BioRender.com/k5ln2gm

### 3.2. Opposing gain- and loss-of-function mechanistic signatures

Network-level summaries of each genotype against WT (Methods §1.5) resolved the direction asymmetry into distinct mechanistic signatures (**Figure 4**). The GoF extreme (SL) was dominated by an inflammatory and tissue-remodeling program: a TGF-β–centred upstream network driving IL-6/STAT3 and IL-1 cytokine signaling, HIF1A and VEGFA, and an extracellular-matrix and matricellular module (S100, SPP1, integrins, collagen and fibrosis signaling). This signature was recovered at both FDR < 0.05 and the stricter FDR < 0.001, indicating that it is not an artefact of threshold choice. A second, independent GoF disease state, heterozygous D/+ on the C3H·B6 (N7) background after ∼20 tonic-clonic seizures at P100, reproduced the same program (**Figure 4B**): the network was organized around TGFB1 and angiotensinogen (AGT) as upstream drivers of an IL-1/IL-6/TNF/IL-17 cytokine hub coupled to NF-κB (MYD88, IKK, RELA) and IL-6/STAT3 signaling, converging on acute-phase response and on hepatic and pulmonary fibrosis. The signature was concordant across normalization schemes (ALL, MATCH) and FDR thresholds (0.05, 0.01). The optimal-dose genotype (LL) was instead characterized by growth-factor and morphogenesis programs (FGF2, VEGFA, BMP4 and TGF-β3, EGF/MET, and RHOA/ROCK– cytoskeletal signaling) that were predominantly decreased, consistent with the global quieting reported above.

**Figure 4.**
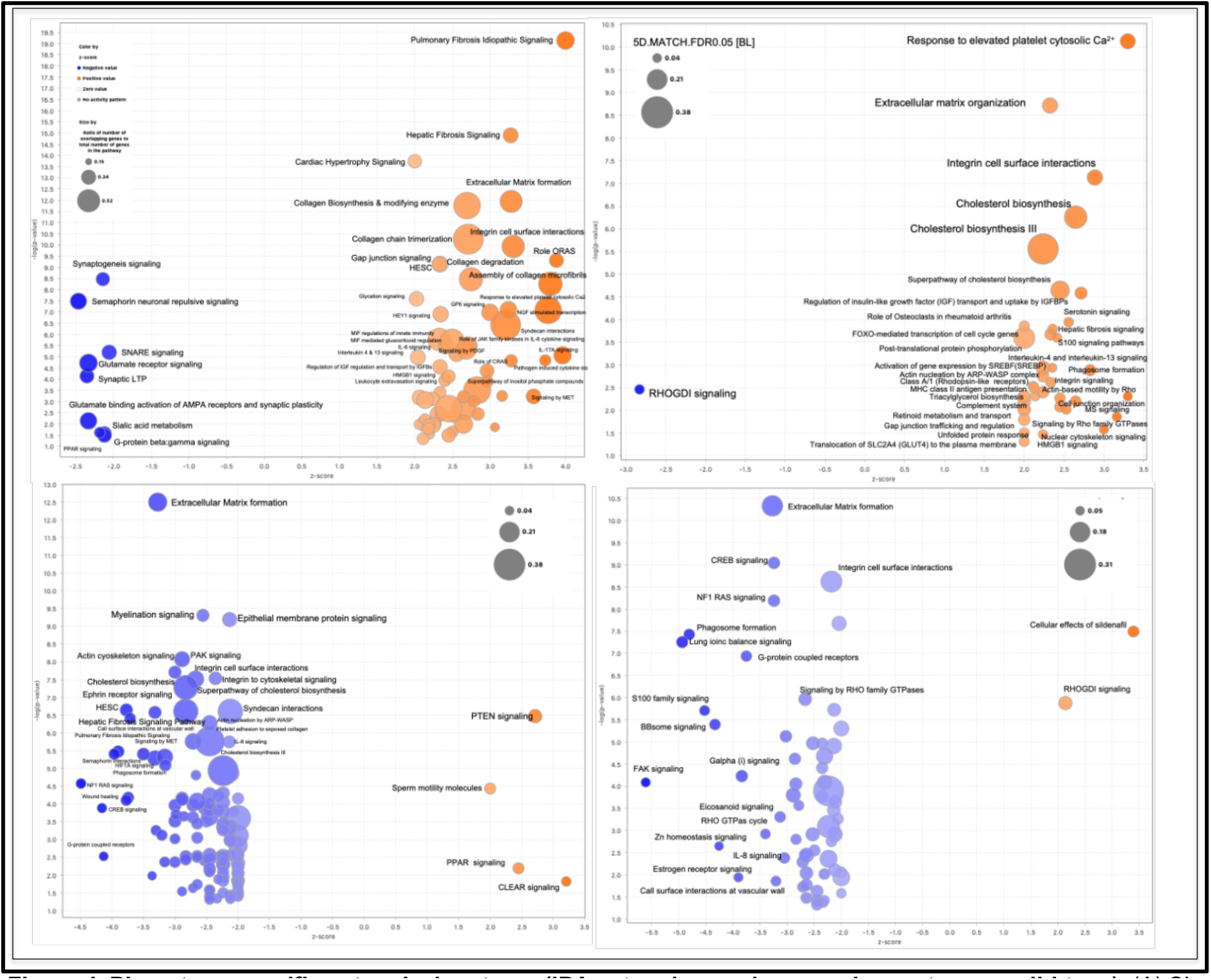
Phenotype-specific network signatures (IPA network overviews, each genotype vs wild-type). (A) SL: a TGF-β–driven inflammatory/IL-6 and extracellular-matrix remodeling signature, shown at FDR < 0.05 and FDR < 0.001. (B) D/+ on the C3H·B6 (N7) background after ∼20 TCs at P100: a TGFB1/AGT-driven inflammatory and fibrotic network (IL-1/IL-6/TNF/IL-17, NF-κB, STAT3) converging on acute-phase response and on hepatic and pulmonary fibrosis, a second GoF disease state reproducing the SL signature. (C) LL: predominantly decreased growth-factor and morphogenesis signaling. (D) HP: suppressed angiogenic and growth-factor signaling with reduced CREB-in-neurons and ion-homeostasis programs. Bubble area, ratio of differentially expressed genes to pathway size; axes, −log10 p-value versus activation z-score. Oncology-derived pathway names are interpreted at the gene/process level (Methods §1.5).

The LoF extreme (HP) presented a third, distinct signature: suppression of angiogenic and growth-factor signaling (VEGFA, EGF/EGFR, FOXM1) together with reduced CREB signaling in neurons and reduced cellular ion-homeostasis programs. The two pathological extremes were thus not graded versions of one program but opposite ones, an active inflammatory and tissue- remodeling signature in GoF disease versus a suppressed growth-factor and neuronal-signaling state in LoF deficiency. This network-level contrast tracked the effect-level annotation of individual pathways (**Figure 4**): pathways such as Hepatic Fibrosis Signaling were activated in SL and deactivated in HP, whereas CREB Signaling in Neurons showed the inverse pattern, with the two inference layers converging on the same gain- /loss-of-function divergence.

### 3.3. A conserved, background-independent gain-of-function injury core

To test whether the injury signature is intrinsic to the GoF state, we compared four z-rich untreated genotypes spanning two backgrounds (B6, C3H·B6), two modifier doses (D+/D+, DL/D+), and two ages (P21, P100). All four disease states were activation-dominated (net activation indices +33 to +155), in contrast to the deactivation-dominated long-lived rescue. A conserved core of 59 injury pathways (**Table S2**) was activated wherever significant across disease states and deactivated in the long-lived genotype: idiopathic pulmonary and hepatic fibrosis, S100, wound-healing, neuroinflammation, cytokine-storm, IL-17A, acute-phase signaling, and an integrin/collagen/syndecan extracellular-matrix module (**Figure 5A**). The core was strongly background-independent at homozygous dose (64% Jaccard overlap of activated injury pathways between B6 and C3H·B6 D/D); at heterozygous dose the response was background-dependent, with C3H·B6 heterozygotes markedly attenuated (**Figure 5B**). Because homozygous samples were necessarily juvenile and heterozygous samples adult, dose and age are confounded; the strong B6-heterozygote signature likely reflects cumulative seizure exposure rather than dose, precluding a clean test of dose-intermediacy. The principal result is robust to this confound: a stereotyped, background-independent GoF injury core is activated wherever disease is expressed and is specifically inverted by the protective modifier.

**Figure 5.**
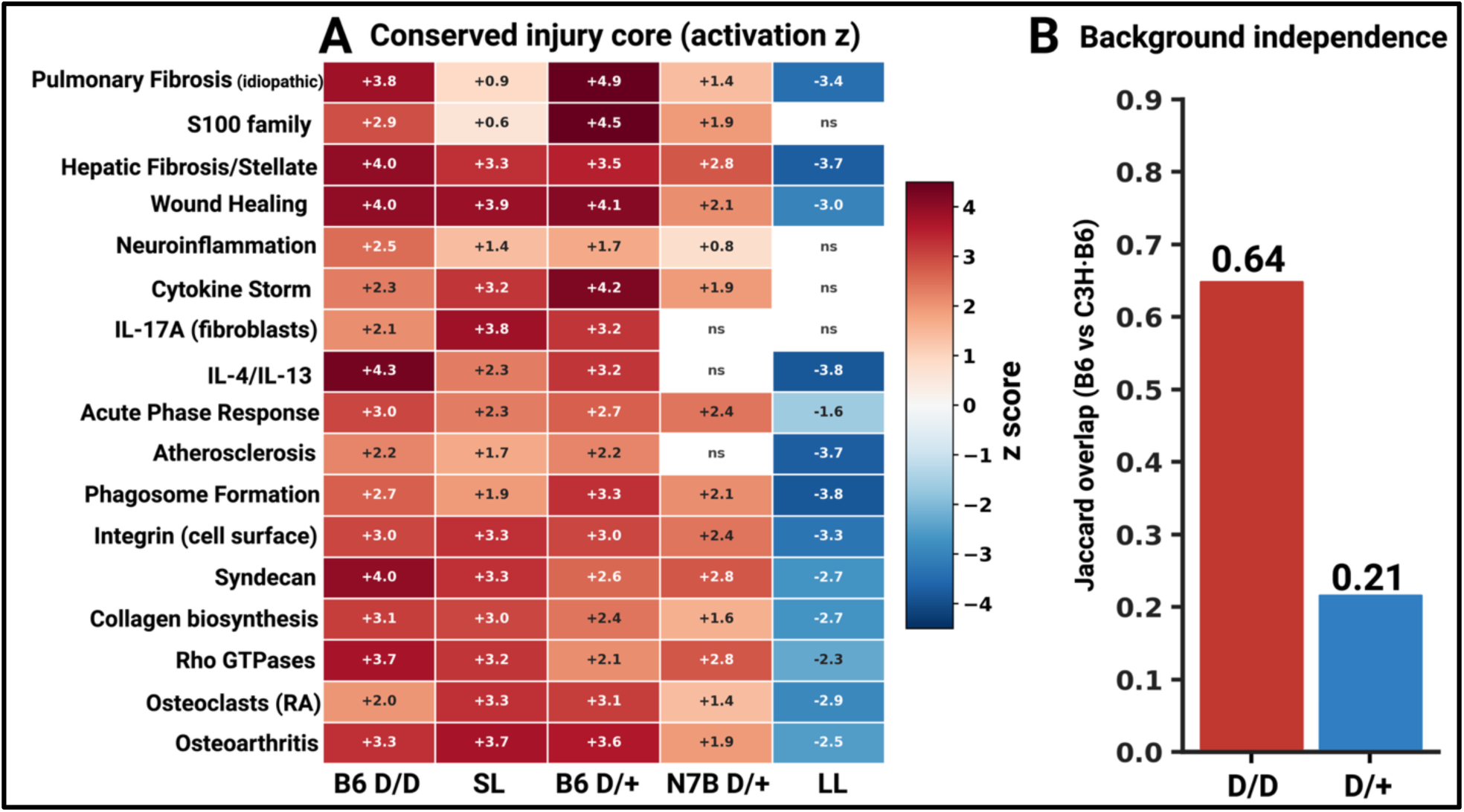
The conserved gain-of-function injury core. (*A*) Activation z-scores for representative core pathways across four disease genotypes and the long-lived rescue; the core is activated wherever disease is expressed (red) and inverted in LL (blue). “ns”, not significant. (*B*) Background concordance (Jaccard overlap of activated injury pathways between B6 and C3H·B6) is high at homozygous dose and attenuated at heterozygous dose. Created in BioRender. Bahramnejad, E. (2026) https://BioRender.com/vvvzsfg.

Extending this comparison to the LoF state placed the injury core across the full dose and direction range (**Figure 4**). The core was deactivated not only in the long-lived rescue genotype but also in HP, often more deeply (for example, idiopathic pulmonary fibrosis and S100 signaling reached activation z of −5.6 and −4.8 in HP). Of the 59 core pathways, 38 were significantly deactivated in HP and only one significantly activated, confirming that the injury core is a feature of the GoF direction specifically and is absent from both the rescue and the LoF states. The LL and HP genotypes were not equivalent, however. They converged in switching the injury core off, but diverged in their wider signatures (**Figure 4**): LL showed quiet deactivation without a neuronal penalty, whereas HP additionally suppressed CREB signaling in neurons and ion-homeostasis programs. The shared deactivation of the injury core therefore coexists with a LoF-specific neuronal signature, a distinction developed in the Discussion.

### 3.4. Durable suppression of the injury cascade in long-lived mice

Within a single long-lived N6 line followed across seizure onset (pre-TC, P42; post-TC, P65), the fibro-inflammatory enrichment signature did not intensify after seizures began. Of 62 pathways enriched pre-TC, 17 (27%) involved injury effects, including fibro-inflammatory core pathways (idiopathic pulmonary fibrosis, JAK/IL-6 signaling); of 36 pathways enriched post-TC, only 3 (8%) involved injury effects, none belonging to the fibro-inflammatory core, with newly enriched pathways instead dominated by metabolic, cell-cycle, and stress-response programs (**Figure S 1**). Activation z-scores were largely unavailable for these contrasts, precluding a directional analysis; this comparison is therefore limited to enrichment presence, and because the only available pre-seizure LL samples occupy markedly different developmental stages (P21 vs P42) we did not compare the pre-seizure transcriptome across ages. With those caveats, the compositional shift away from the injury core argues against intensification of the cascade after seizure onset.

### 3.5 Convergence of genetic and pharmacologic rescue on a shared injury core

To ask whether the protective modifier and the disease-modifying drug candesartan act on the same downstream program, we compared the conserved injury core across untreated, candesartan-treated, and modifier-bearing genotypes (**Figure 6**). In untreated B6 D/D hippocampus all 59 core pathways were significantly enriched; candesartan reduced this to 1 of 59, mirroring a global collapse of the disease transcriptome (488 → 28 significantly altered pathways) — normalization toward the WT baseline (**Figure 6A,B**). In the longer-lived heterozygous arm candesartan removed the core partially (14–28 of 59 remaining across treatment timepoints). The genetic modifier produced a distinct pattern: in long-lived hippocampus 48 of 59 core pathways remained significantly altered, but 40 were deactivated (i.e., the core was inverted rather than removed) and the overall footprint remained large (313 pathways) and distinct from WT. Despite these divergent global strategies (i.e., pharmacologic normalization toward baseline versus genetic suppression below it), the interventions converged on a shared target program: 39 conserved injury-core pathways were both removed by candesartan and inverted by the modifier (**Figure 6C**), including fibrosis, acute-phase, atherosclerosis, leukocyte-adhesion, collagen, and cytoskeletal-remodeling signaling. Genetic reduction of Na_V_1.6 and pharmacologic AT1R blockade thus neutralize the same seizure-associated injury cascade by mechanistically distinct routes.

**Figure 6.**
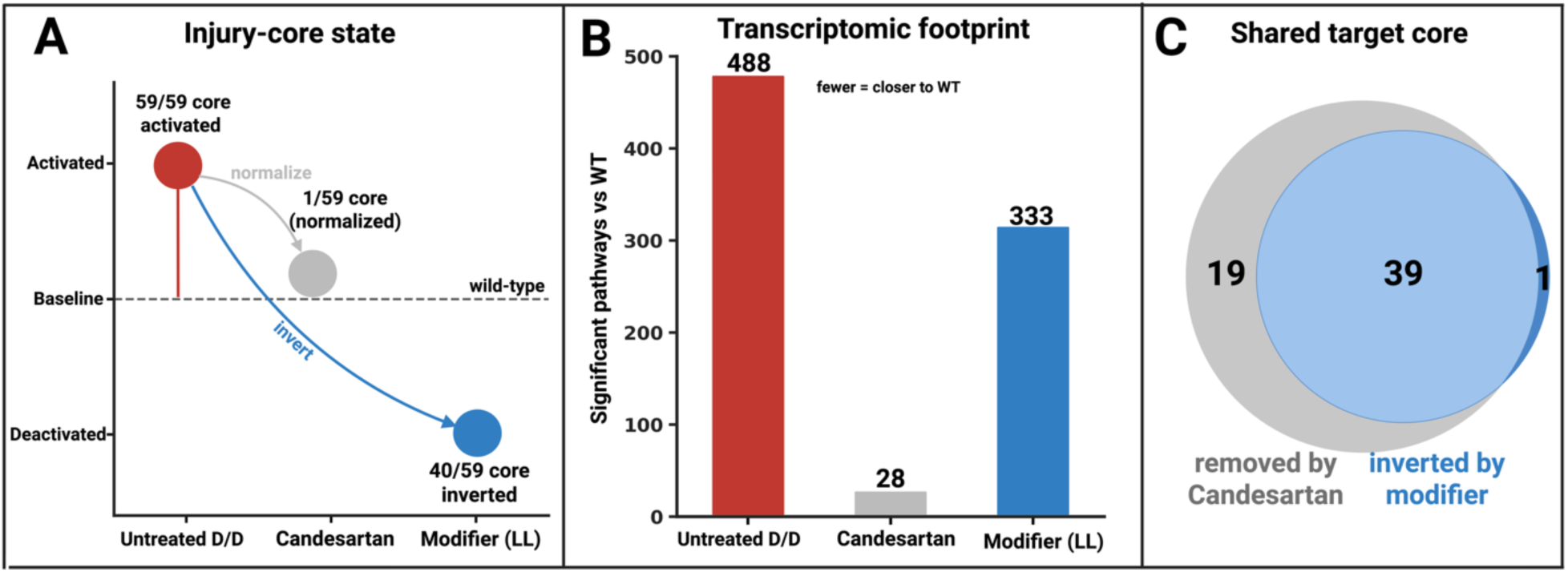
Two routes to rescue of the *Scn8a*-N1768D injury cascade. (*A*) State of the conserved injury core relative to WT: activated in untreated D/D (59/59 enriched), normalized by candesartan (1/59), inverted by the modifier (40/59 deactivated). (*B*) Transcriptomic footprint (significantly altered pathways vs WT): candesartan collapses the disease footprint toward WT (488→28); the modifier maintains a large, distinct footprint (313). (*C*) Of the conserved core, 39 pathways are both removed by candesartan and inverted by the modifier — a shared target program reached by distinct routes. Created in BioRender. Bahramnejad, E. (2026) https://BioRender.com/45ufxwf.

## 4. Discussion

Taken together, these analyses describe how a single *cis*-acting *Scn8a* modifier reshapes the hippocampal transcriptome of N1768D homozygotes in a dose-dependent manner. As modifier copy number rises and Na_V_1.6 expression falls (SL > LL > HP), the transcriptome shifts from florid global activation (SL) through global deactivation at the protective dose (LL) to a further- deactivated, LoF state (HP). The protective optimum is therefore not a selective correction of disease pathways but a globally low-tone transcriptional state, consistent with a Na_V_1.6 dose- response in which both excess and deficiency are pathological and an intermediate level is protective (“Goldilocks Zone”) (**Figure 7**) (10).

**Figure 7.**
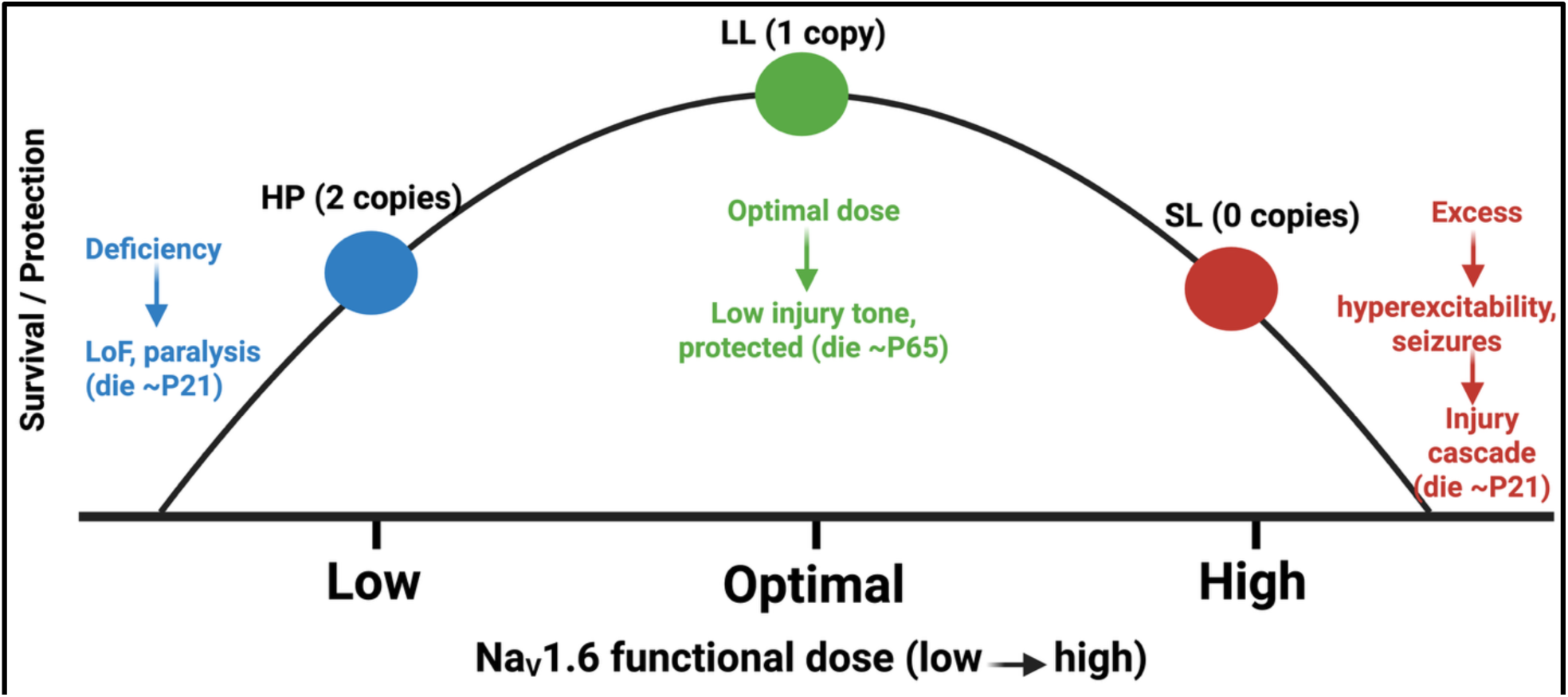
Na_V_1.6 dose-response model. Both excess (SL, no modifier copies) and deficiency (HP, two copies) of Na_V_1.6 are lethal in the juvenile period (SL ∼P21, HP ∼P21); an intermediate modifier dose (LL, one copy) is protective (survival ∼P65). Excess drives hyperexcitability, seizures, and the injury cascade; deficiency causes loss of function and paralysis. Created in BioRender. Bahramnejad, E. (2026) https://BioRender.com/wv2re0v.

### 4.1 Opposing gain- and loss-of-function transcriptional programs

The most informative feature of the series is that its two lethal extremes are mechanistically opposite rather than two degrees of one process. The GoF extreme (SL) engages an inflammatory and tissue-remodeling program (TGF-β, IL-6/STAT3, HIF1A and VEGFA, and an extracellular-matrix module), the molecular correlate of the seizure-driven injury cascade considered below. The LoF extreme (HP) shows the converse: suppression of growth-factor and angiogenic signaling together with reduced CREB signaling in neurons and disturbed ion homeostasis, a signature of perturbed neuronal signaling and arrested growth rather than of active injury. This within-animal divergence parallels the clinical dichotomy of human SCN8A disease, in which GoF variants cause seizure-dominated developmental and epileptic encephalopathy and LoF variants more often cause intellectual disability, movement disorder, or absence epilepsy with a distinct comorbidity profile (7, 10, 24, 25). To our knowledge no prior study has contrasted GoF and LoF hippocampal transcriptional mechanisms within a single genetic model, and the parallel suggests that the contrasting human presentations have, at least in part, distinct tissue-level molecular correlates. The same dichotomy is established in the mouse. Severe loss of Na_V_1.6, as in the *Scn8a*-null (med) allele, produces dystonia and perinatal lethality rather than seizures, and hypomorphic alleles such as V929F and A1071T produce ataxia with absence seizures, whereas GoF alleles (N1768D, R1872W, T767I) produce convulsive seizures and premature death (26, 27). The HP transcriptome, with its suppression of neuronal CREB and ion-homeostasis programs and its lack of the injury core, is consistent with this LoF pole.

### 4.2 Loss-of-function mechanisms and the human disorder

The HP signature offers a tissue-level account of why LoF produces a phenotype so different from GoF. Rather than the inflammatory and remodeling cascade that dominates the seizing brain, the LoF state is defined by suppression of the programs that build and tune neurons: CREB signaling in neurons, ion homeostasis, and growth-factor and angiogenic signaling. CREB-dependent transcription is a convergent node for synaptic plasticity, learning, and memory, so its suppression provides a plausible molecular correlate for the intellectual disability and developmental delay that characterize human LoF *SCN8A* variants, while the ion-homeostasis and growth-factor deficits align with the movement and developmental features (16, 28, 29). The implication is that LoF disease is not a milder version of the GoF injury process but a mechanistically distinct, deficit- driven state. This carries a direct therapeutic corollary: the two directions call for opposite logic. GoF pathology is addressed by lowering channel activity or its downstream cascade, through antisense reduction of *Scn8a*, isoform-selective block, or anti-inflammatory AT1-receptor blockade, whereas LoF pathology would instead require restoring functional channel output, for example by enhancing expression of the functional allele (11, 26, 30). Interventions that benefit the GoF state, including dose reduction and anti-inflammatory treatment, would be expected to be ineffective or harmful in the LoF state.

A second feature of the series clarifies how the HP genotype maps onto the human LoF spectrum. Although HP hippocampus retains roughly 30% of WT *Scn8a* transcript, its phenotype, hindlimb paralysis and euthanasia by ∼P21, is that of a near-complete null rather than a partial hypomorph. The resolution lies in the distinction between transcript abundance and functional channel output. The modifier is a *de novo* 2-bp deletion/frameshift mutation in exon 15, which encodes the DIIS4 voltage sensor; the transcript that escapes nonsense-mediated decay is therefore translated only as far as domain II, truncating Na_V_1.6 before the domain III and IV pore-forming segments and before the N1768D residue itself, and an independent exon-15 frameshift allele is likewise predicted to be null (31). The residual mRNA thus yields no functional channel, so in homozygotes (HP) functional Na_V_1.6 approaches zero despite ∼30% residual transcript. This places HP at the severe-deficiency pole occupied by the *Scn8a*-null (med) allele, which causes hindlimb paralysis and juvenile lethality (27), rather than at the milder hypomorphic pole of *Scn8a*-medJ (5–10% expression), whose homozygotes survive to adulthood with ataxia and dystonia. The same truncation explains the modifier’s protective action in cis: by preventing translation of the N1768D GoF residue, a single copy removes the mutant channel from one chromosome (LL) without abolishing Na_V_1.6.

This functional-null interpretation also reframes the cross-species comparison. The phenotypic consequence of reduced Na_V_1.6 differs between mouse and human chiefly because the two species occupy different points on the haploinsufficiency curve. Mice tolerate a 50% reduction with only subtle consequences, heterozygous *Scn8a*-null mice are largely normal but show spike-wave discharges and mild neurobehavioral changes (32), whereas the equivalent loss in humans produces intellectual disability and autism spectrum features, with or without (typically absence) seizures (29, 33–35). Because mice are not haploinsufficient for *Scn8a*, a comparable functional-loss threshold is reached by different genetic routes in the two species: in mice through homozygous hypomorphic or null alleles (or, here, a homozygous functional-null modifier), and in humans through a single heterozygous variant that severely reduces channel activity and, in some cases, may act dominantly on the WT allele. Such a dominant mechanism has been proposed for

Na_V_1.6 through α-subunit dimerization (36, 37) and is well established for the paralogous cardiac channel Na_V_1.5 (38), but it has not been directly demonstrated for Na_V_1.6 and remains a plausible extension. Across this range, human severe loss of function presents with movement disorder, from isolated myoclonus without seizures or cognitive impairment to profound hypotonia (35, 37), the pole that HP models. The apparent species difference in severity therefore reflects differing haploinsufficiency thresholds and genetic architecture, not a fundamentally more severe murine response to comparable functional loss.

### 4.3 A conserved, seizure-driven injury core and its durable rescue

The disease signature itself proved remarkably stereotyped. A conserved core of injury pathways consisting of fibro-inflammatory programs with an extracellular-matrix and cytoskeletal module was activated in every GoF genotype examined and was strongly background-independent at homozygous dose. Because this core was activated in post-seizure animals but absent or deactivated in pre- and non-seizing genotypes of the same Na_V_1.6 dose, it behaves as a seizure-associated injury cascade rather than a direct readout of channel dosage, bearing on the channel- versus-cascade question and indicating that the modifier’s protection operates upstream of the cascade by limiting the seizure burden that drives it.

Within the long-lived genotype, the cascade did not intensify after seizure onset; if anything its fibro-inflammatory enrichment diminished, with the post-seizure transcriptome shifting toward metabolic and stress-response programs. For instance, Nerve Growth Factor (NGF)-stimulated transcription signaling was the single most activated pathway in both pre-seizure (P42) (-log_10_ p-value= 16.9, z-score= 3.0) and post-seizure (P65) (-log_10_ p-value= 7.1, z-score= 2.0) N6 D/D mice (i.e., carrying one copy of the linked modifier). Activating the NGF-stimulating transcription pathway is critical in brain injury because it promotes neuronal survival, triggers axonal regeneration, and reduces neuroinflammation. By bypassing cell death programs, this pathway restores lost cognitive and motor functions following trauma (39). Although this comparison was limited to enrichment presence and to a single line at a later developmental window, it suggests that the modifier confers durable rather than transient protection: by holding Na_V_1.6 and seizure burden low, it appears to prevent the feed-forward injury cascade from becoming self-sustaining, rather than merely delaying it.

### 4.4 Two routes to one target: genetic rescue and a repurposed drug

Comparison with our pharmacologic candesartan data revealed both a convergence and an instructive divergence (**Figure 6**). Genetic dose-reduction and AT1R blockade act on the same conserved injury core (i.e., 39 core pathways were both removed by candesartan and inverted by the modifier), identifying this fibro-inflammatory cascade as a central, drug-accessible node in *Scn8a* GoF pathology, reachable either by lowering channel dose or by blocking angiotensin signaling. Yet the two interventions neutralize it by opposite transcriptomic strategies: candesartan normalizes the transcriptome toward the WT baseline (the disease footprint collapsing from 488 to 28 altered pathways, and the injury core from 59/59 enriched to 1/59), whereas the modifier suppresses the core below baseline while maintaining a large, distinct footprint (313 altered pathways; 40/59 core pathways inverted). A disease-modifying drug restores normalcy, while the protective allele installs a different, quieted state that happens to disarm the same cascade. The present results add an upstream rationale for this convergence. The heterozygous GoF disease network (N7 D/+) was organized around angiotensinogen (AGT) as a predicted upstream regulator (**Figure 4B**), indicating engagement of the brain renin-angiotensin system in the seizing hippocampus. Because candesartan acts on this same axis through AT1-receptor blockade, the genetic and pharmacologic routes converge not only at the injury core but at an upstream node: GoF disease activates angiotensin signaling, and candesartan interrupts it. This links the present series to our prior demonstration that candesartan restores blood-brain-barrier integrity, normalizes aberrant gene expression, and extends survival in the same model [(17), and to the broader role of brain renin-angiotensin signaling in epileptogenesis (2). AT1-receptor overstimulation is known to drive NF-κB-dependent production of IL-1β, TNF, and IL-6 and, in models of brain injury, to promote TGF-β-driven fibrosis, the same nodes that dominate the N7 D/+ disease network, so AT1-receptor blockade is positioned to act on the GoF injury program itself rather than on channel activity.

### 4.5 A shared injury program across neurological disease

The GoF injury signature is not specific to *SCN8A*. Processed through the same differential-expression and pathway-inference pipeline, a humanized α-synuclein mouse model of Parkinson’s disease (40) and hippocampal tissue from temporal lobe epilepsy patients with low seizure frequency (41) were both activation-dominated (net activation indices of +60 and +83), like the GoF states of the present series. A substantial fraction of the conserved injury core was activated in each (23 of 59 core pathways in the Parkinson’s model, 16 of 59 in low-frequency epilepsy), and 75% of the core pathways activated in SL were also activated in the Parkinson’s model. In both, the shared program was the same IL-1/IL-6/TNF cytokine module, wound-healing signaling, and hepatic-fibrosis and stellate-cell activation seen in the mouse, and angiotensinogen again appeared as an upstream driver in the Parkinson’s network, reinforcing the renin-angiotensin link noted above. The GoF injury core is therefore better understood as a general transcriptional response to chronic CNS injury than as an *SCN8A*-private one (22).

Human epilepsy at high seizure frequency behaved differently. The high-frequency cohort was deactivation-dominated (30 activated versus 75 deactivated pathways), the injury core was largely absent, and neuronal-system programs (calcium, CREB in neurons, and opioid signaling) were systematically suppressed. As argued previously, this widespread deactivation at the severe end of seizure burden is most parsimoniously explained by neuronal loss rather than by coordinated regulation across many pathways, paralleling the synaptic-gene downregulation attributed to neuron and synapse loss in Alzheimer’s disease (41, 42). This comparison also sharpens an interpretive point for the present series: a deactivated transcriptional readout is not self-explaining. The same downward signal marks a protective inversion in the long-lived genotype, a LoF deficit in HP, and tissue loss in high-frequency human epilepsy, so direction alone does not identify mechanism, and the genotype-specific priors applied here are what separate these cases. The series models injury direction and shares program-level features with these conditions without phenocopying any of them.

### 4.6 Limitations and conclusions

Several cautions bound this interpretation. HP models the severe-deficiency pole of LoF. Its paralysis and respiratory failure resemble the near-null *Scn8a* phenotype (27) more than the milder LoF variants seen in many patients. Thus, the mapping to the human LoF spectrum is directional and conceptual (i.e., the mouse is a model of the human condition, not a phenocopy). A parallel caveat applies at the GoF pole. Mice can show more severe outcomes than humans carrying equivalent variants: whereas children with severe GoF variants rarely die from convulsive status epilepticus, the same variants (e.g., N1768D) are consistently fatal in murine models (5). This discrepancy may reflect differences in brain scale, since in the much smaller murine brain seizure activity may propagate more readily into subcortical structures such as the brainstem, driving sudden death. The series therefore captures the direction and tissue-level mechanism of each pole rather than the human clinical severity at either. The signatures are themselves inferential: derived from network-level summarization and literature-based effect annotation, they are hypothesis-generating rather than hypothesis-testing and await biological validation. Their value is nonetheless strengthened by the convergence of two independent inference layers. The network overview and the beneficial/detrimental annotation independently recover HP’s deactivation of CREB-in-neurons and SL’s activation of fibrosis signaling, and by the direction asymmetry, which is annotation-independent. We note that the pathways carrying oncology-derived names were read at the level of their constituent immune, inflammatory, remodeling, and growth-factor genes, which reflects the shared machinery of tissue scarring, angiogenesis, and growth.

These results also bear on the analytic framework. The beneficial/detrimental pathway-effect inference used here, re-derived under current methods, was concordant with our prior published annotations in 98.6% of co-annotated calls and now connects three independent datasets: the candesartan mouse model, human temporal-lobe-epilepsy resections, and the present modifier series. The human TLE study cautioned that the valence of inflammation is context-dependent rather than fixed (22), and our deliberately context-explicit encoding, together with the genotype-specific scoring, was required precisely because the modifier genotypes occupy different disease states. Two limitations temper the interpretive layer: the beneficial/detrimental tallies depend on those genotype-specific priors (the underlying direction asymmetry does not), and several comparisons are constrained by structural confounds: dose with age (homozygotes are necessarily juvenile) and the absence of activation z-scores in the treated and longitudinal contrasts. A broader literature contextualization, and biological validation of the inferred effects, remain for subsequent work.

In sum, an isogenic *Scn8a* allelic series resolves two questions that single-genotype models cannot. First, the gain- and loss-of-function extremes are mechanistically opposite, an active fibro- inflammatory injury program versus a deficit in neuronal-signaling and growth programs, so the two clinical directions of *SCN8A* disease have distinct tissue-level correlates and call for opposite therapeutic logic. Second, an intermediate channel dose installs a durable, low-tone protective state that converges with a repurposable drug on the same injury core, identifying that core as a tractable target. Because the same injury program is engaged in a Parkinson’s model (40) and in human epileptic tissue, the mechanisms defined here are likely to extend beyond *SCN8A* to the broader problem of seizure- and injury-driven neurological disease.

## Clinical Perspectives

• SCN8A-lowering therapies are advancing toward the clinic, yet it remains unknown what lowering Na_V_1.6 does to the brain beyond suppressing seizures, or whether excessive lowering carries a distinct molecular cost; we therefore compared hippocampal transcriptomes across an isogenic allelic series in which zero, one, or two copies of a cis-acting modifier titrate Scn8a expression.

• Seizing D/D mice activated a coordinated fibro-inflammatory injury program spanning TGF-β, IL-6/STAT3, and extracellular matrix remodelling that recurred across independent genetic backgrounds and overlapped signatures reported in human temporal lobe epilepsy; a single modifier copy (approximately 30% transcript reduction) inverted this program and rescued the epilepsy, whereas two copies (approximately 70% reduction) also silenced it but instead deactivated neuronal growth-factor, CREB, and synaptic signalling, yielding paralysis without seizures.

• Because the two lethal extremes engage opposite transcriptional programs, the injury cascade supplies candidate pathway-level biomarkers for monitoring the efficacy of SCN8A-lowering therapy while the suppressed growth and synaptic signature marks over-lowering, together providing a molecular readout of the therapeutic window and nominating seizure-driven injury pathways, including angiotensin and TGF-β signalling, as adjunctive targets in SCN8A-related developmental and epileptic encephalopathy.

## Data Availability

The data that support the findings of the present study are available from the corresponding author on reasonable request. RNA-sequencing data have been submitted to the Figshare repository at https://doi.org/10.6084/m9.figshare.33023150.

## CRediT Author Contribution

**Michael F. Hammer:** Conceptualization, Resources, Writing—original draft, Funding acquisition, Methodology, Validation.

**Erfan Bahramnejad:** Conceptualization, Data curation, Formal analysis, Methodology, Visualization, Investigation, Project administration, Writing—review & editing.

## Ethics statement

Animal work was approved by the University of Arizona IACUC (protocol #16-160), performed in ARRIVE-compliant accredited facilities, and mice were euthanized at endpoint by cervical dislocation.

## Acknowledgment

This study was primarily supported by the Shay Emma Hammer Research Foundation. RNA- seq was supported by a Core Facility Pilot Program award from the Office of Research and Partnerships at the University of Arizona. We thank the Arizona Genetics Core and the Arizona Genomics Institute for assistance with RNA-seq and whole-genome sequencing, respectively. We also thank Taylor Camacho for assistance with animal toe clipping and genotyping.

## Supplementary Material

**Table S1.**
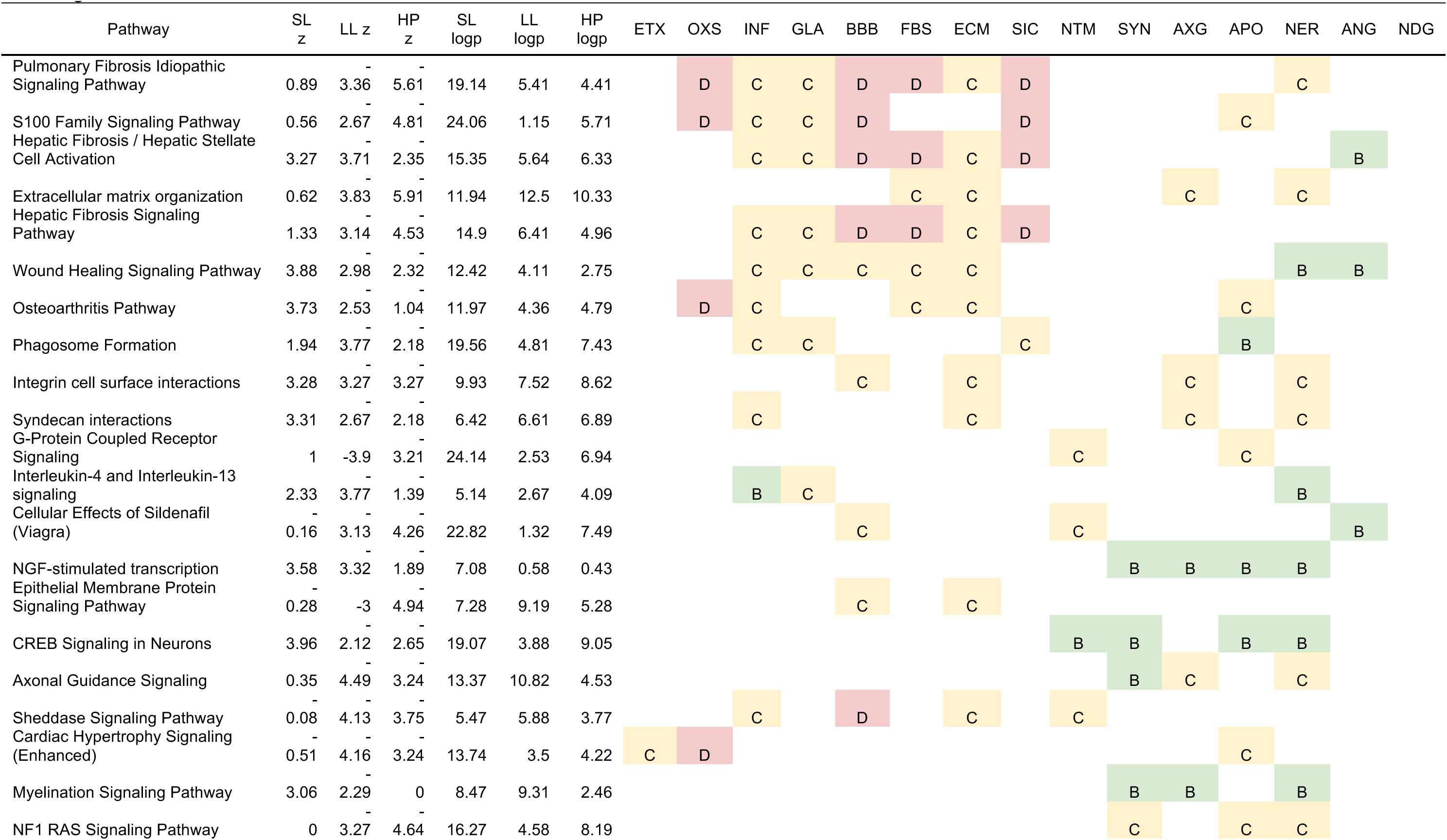
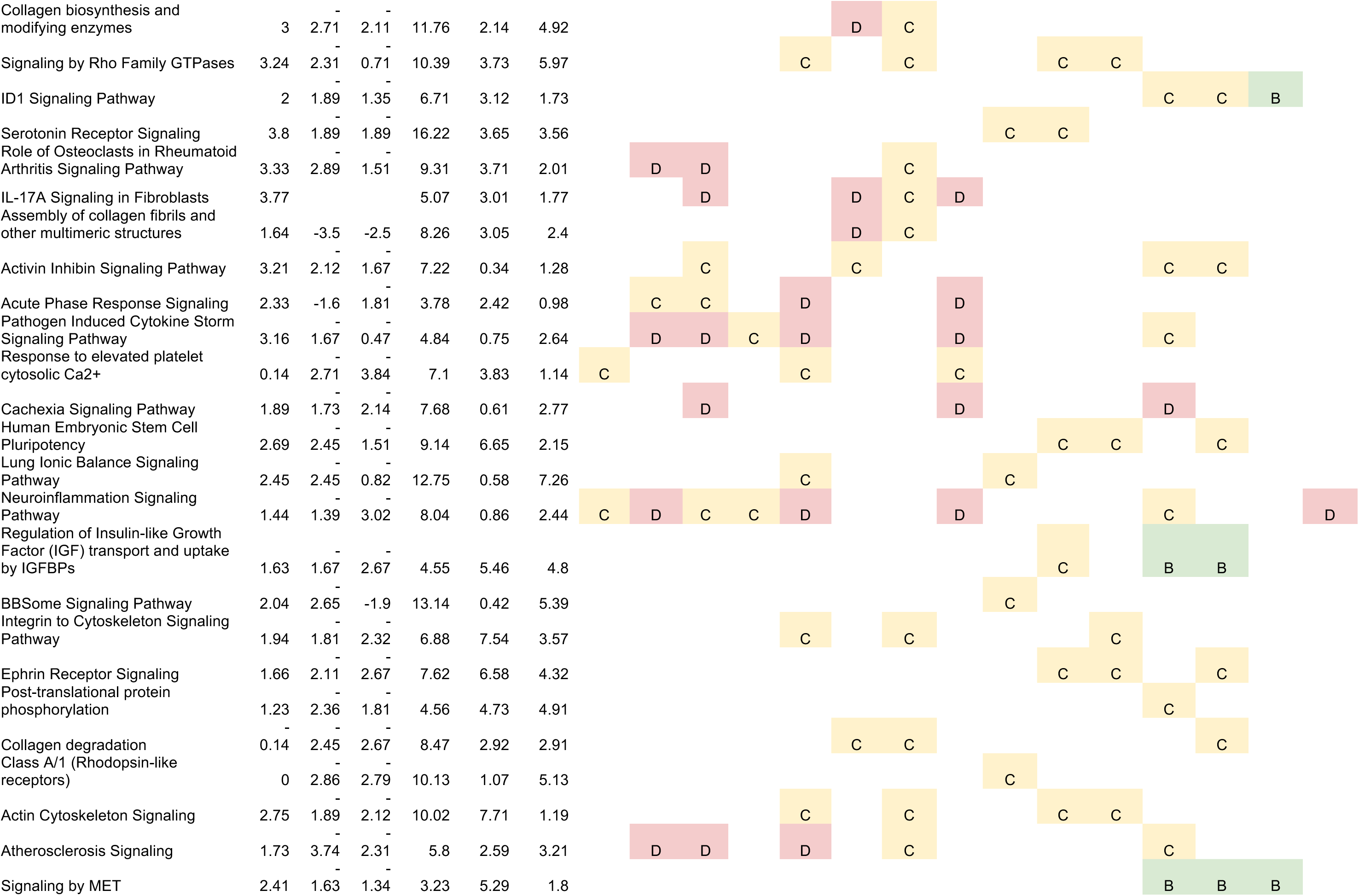
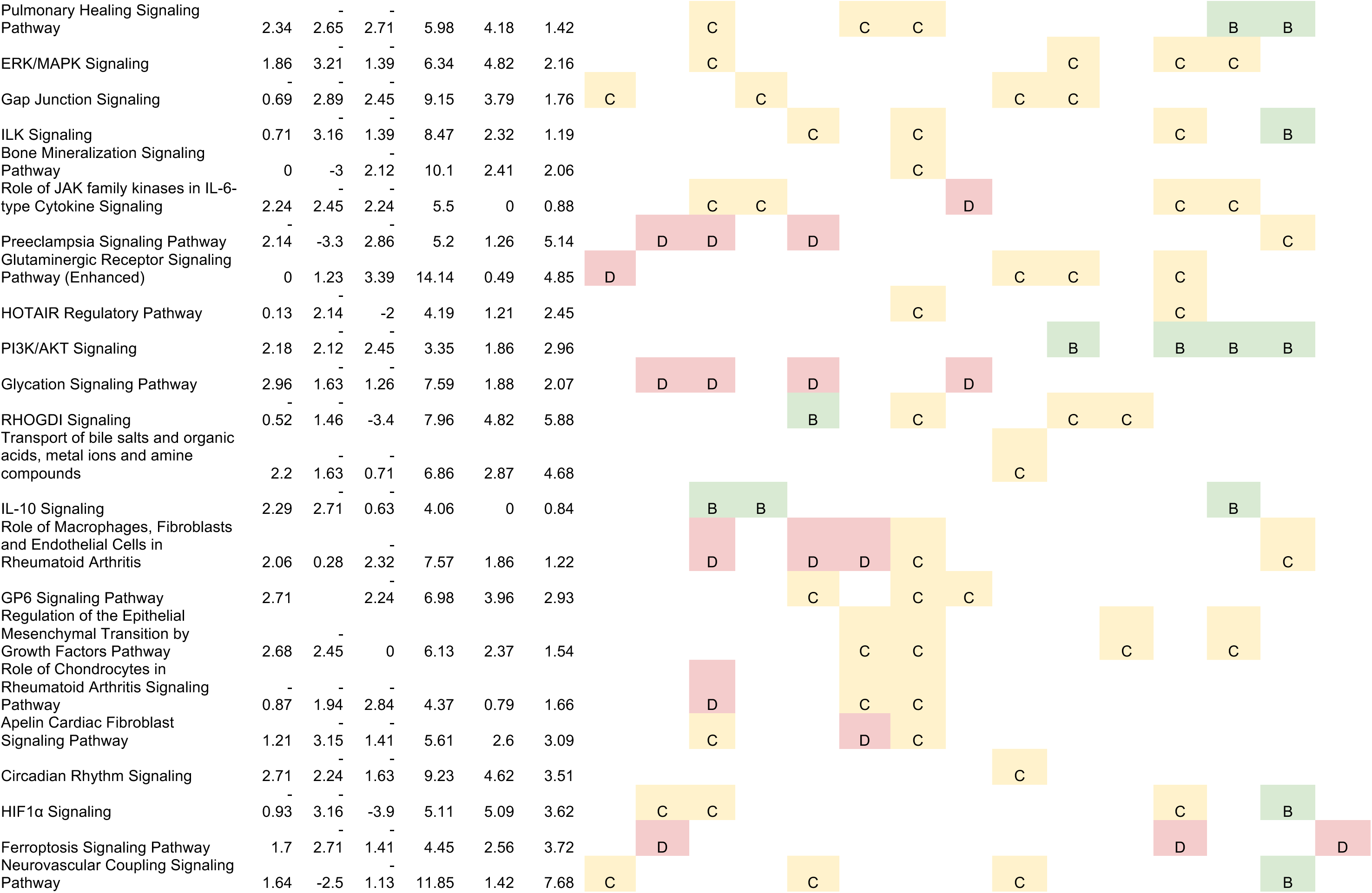
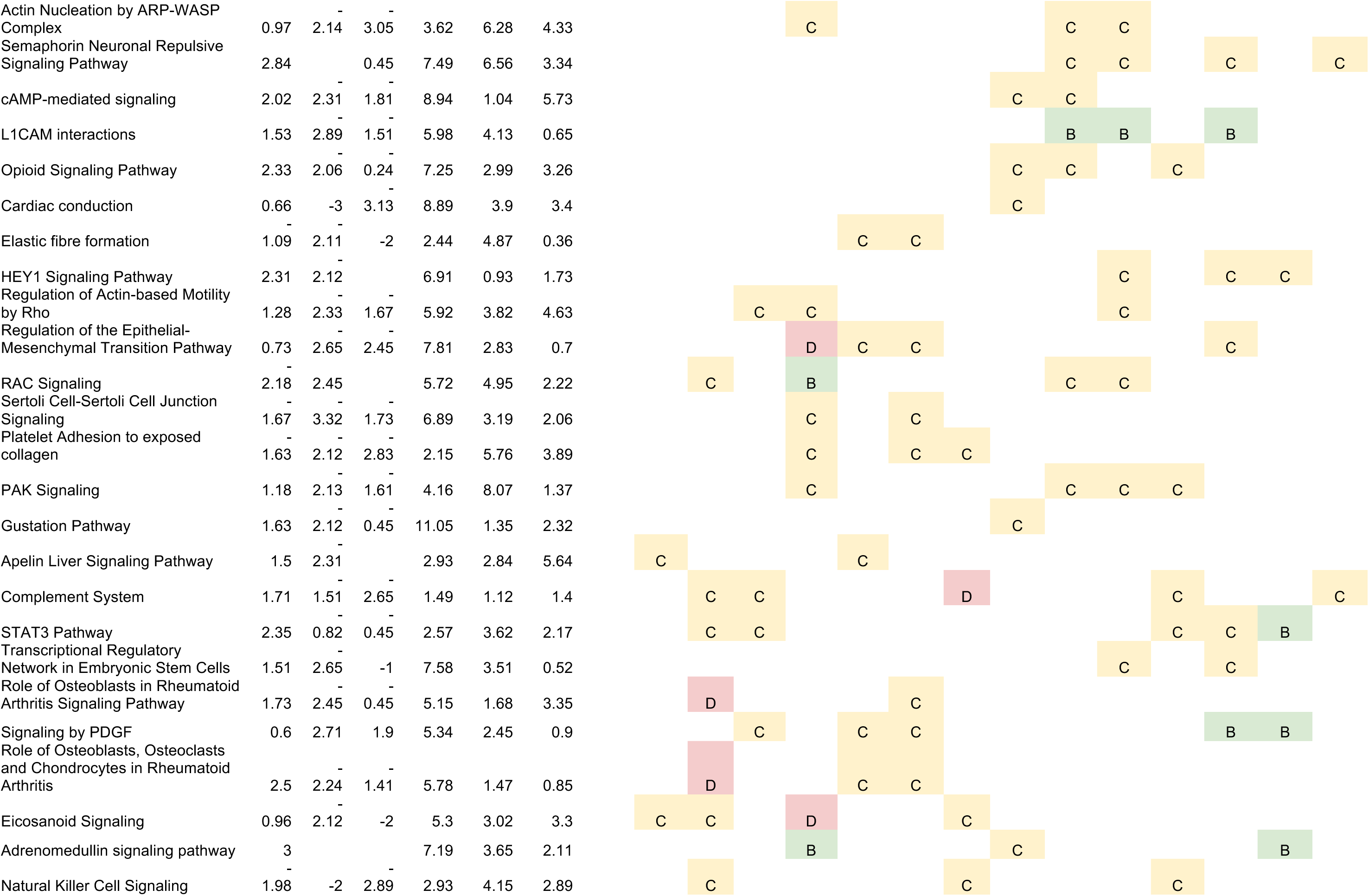
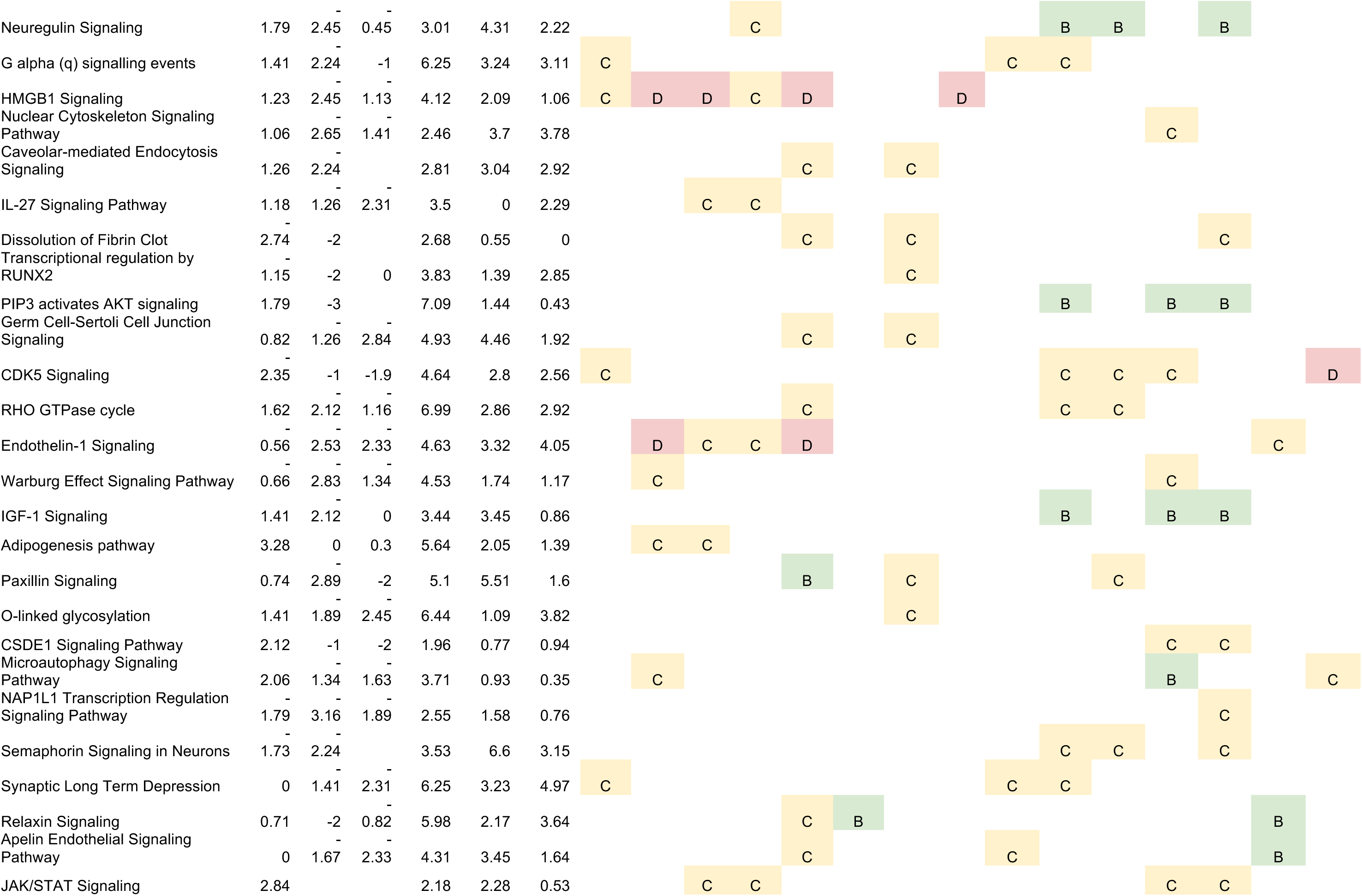
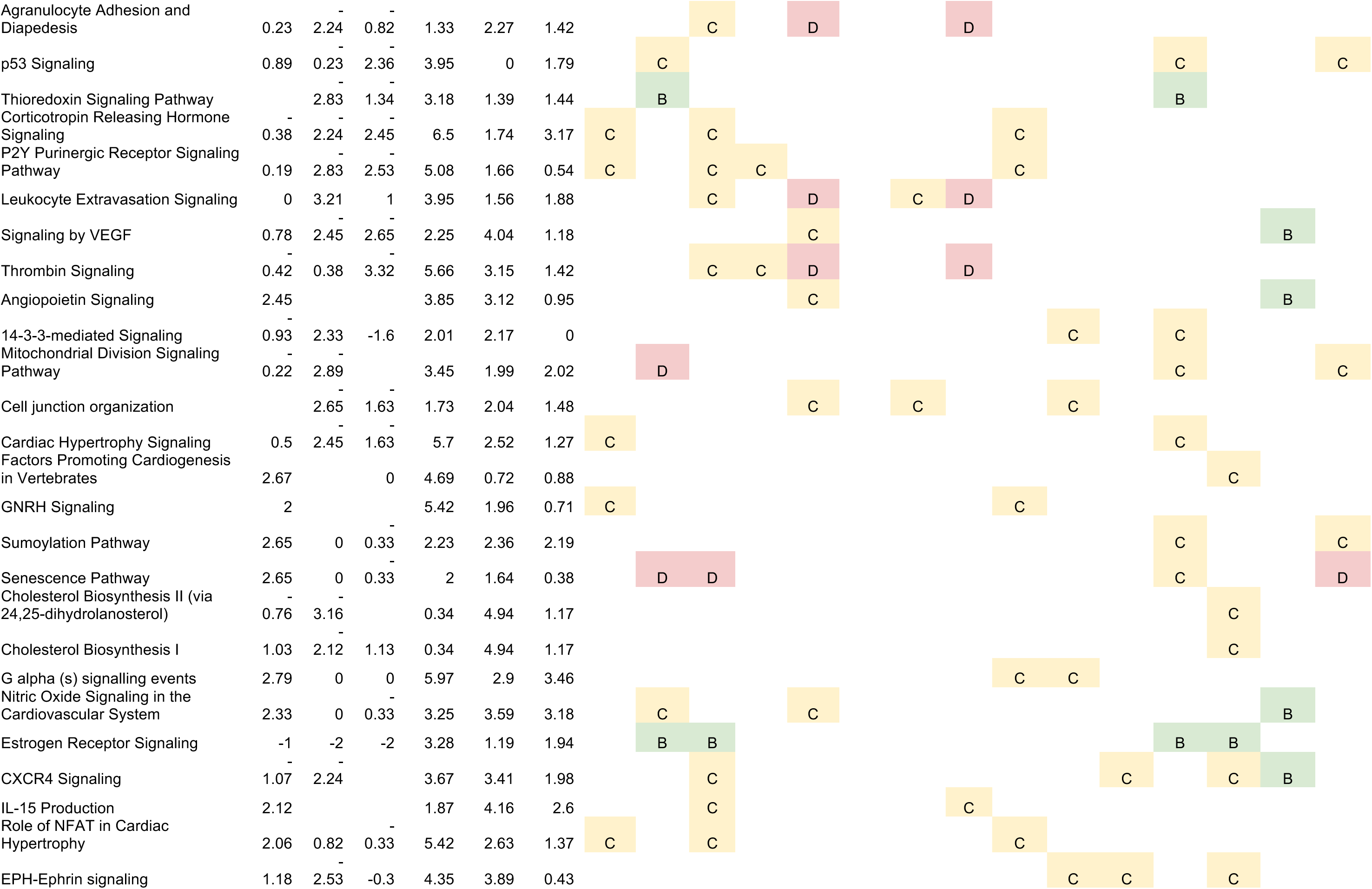
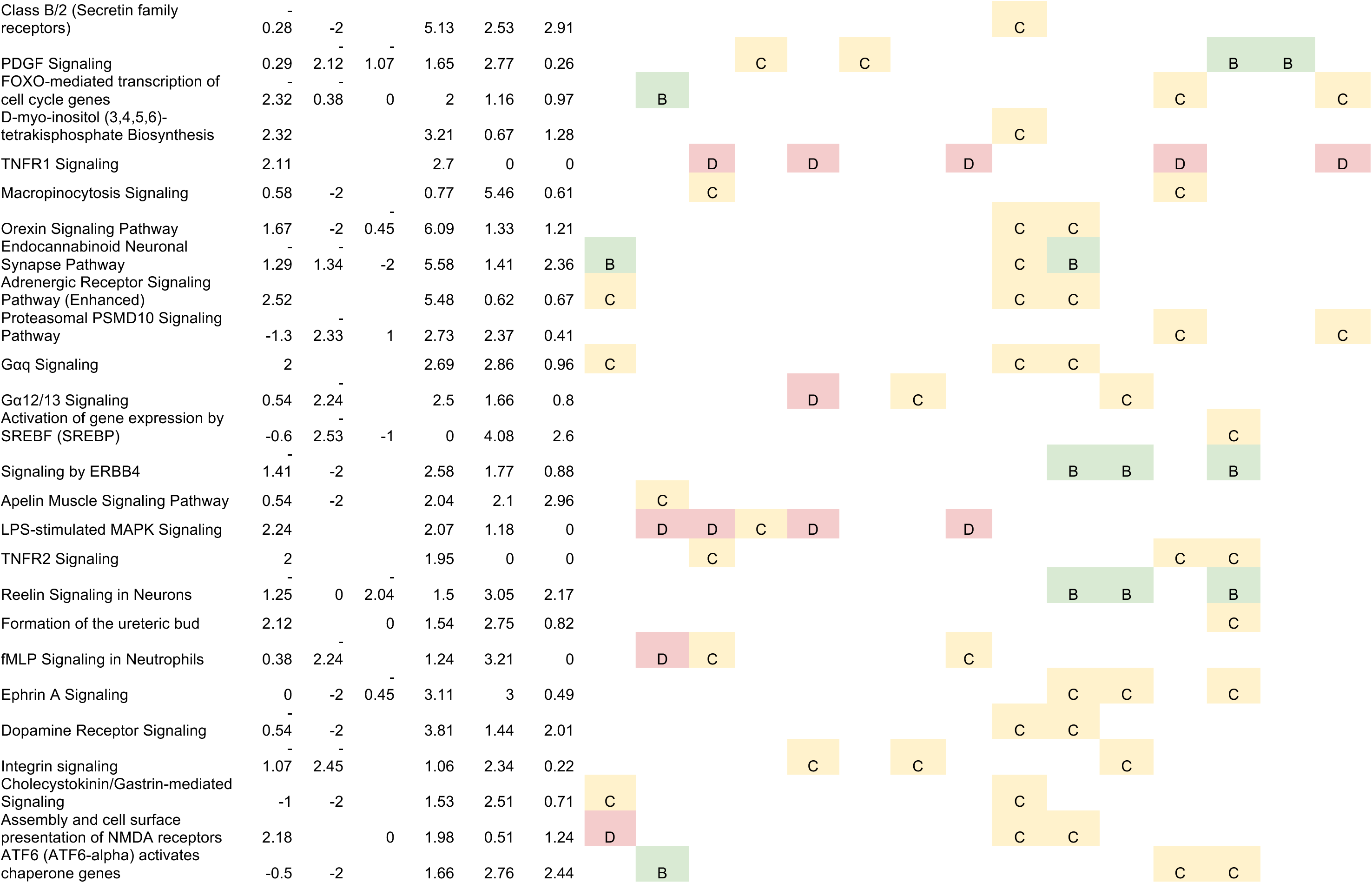
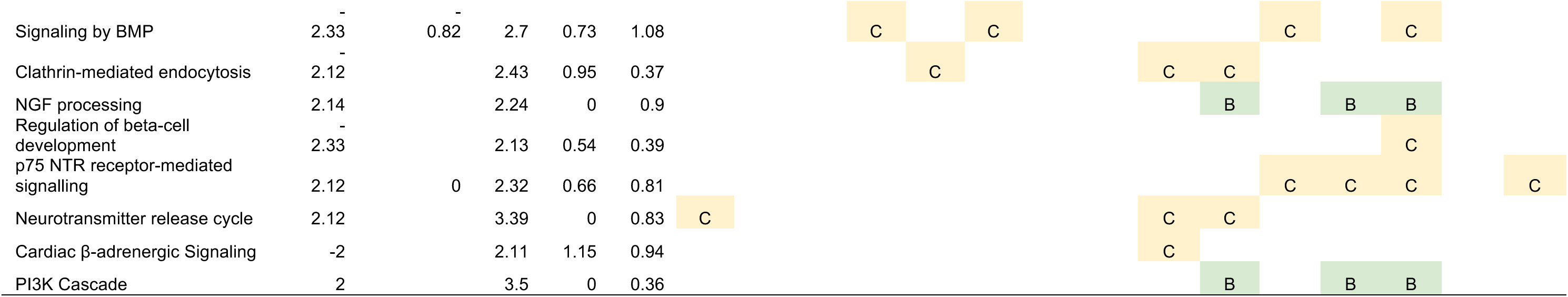
Activation z-scores, enrichment significance, effect annotations, and annotation confidence for the 180 canonical pathways significant and directional in at least one of SL, LL, or HP.

**Table S2.**
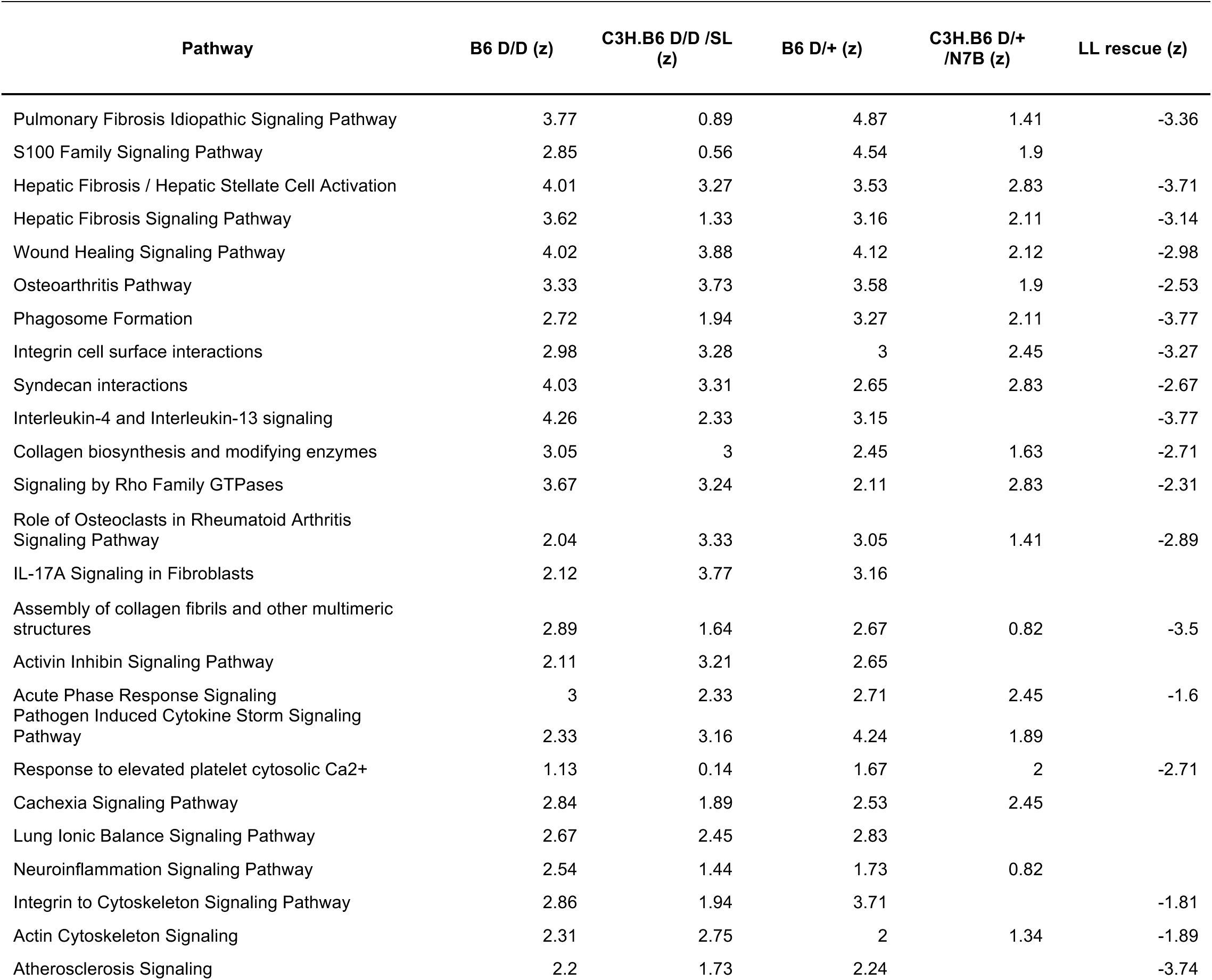
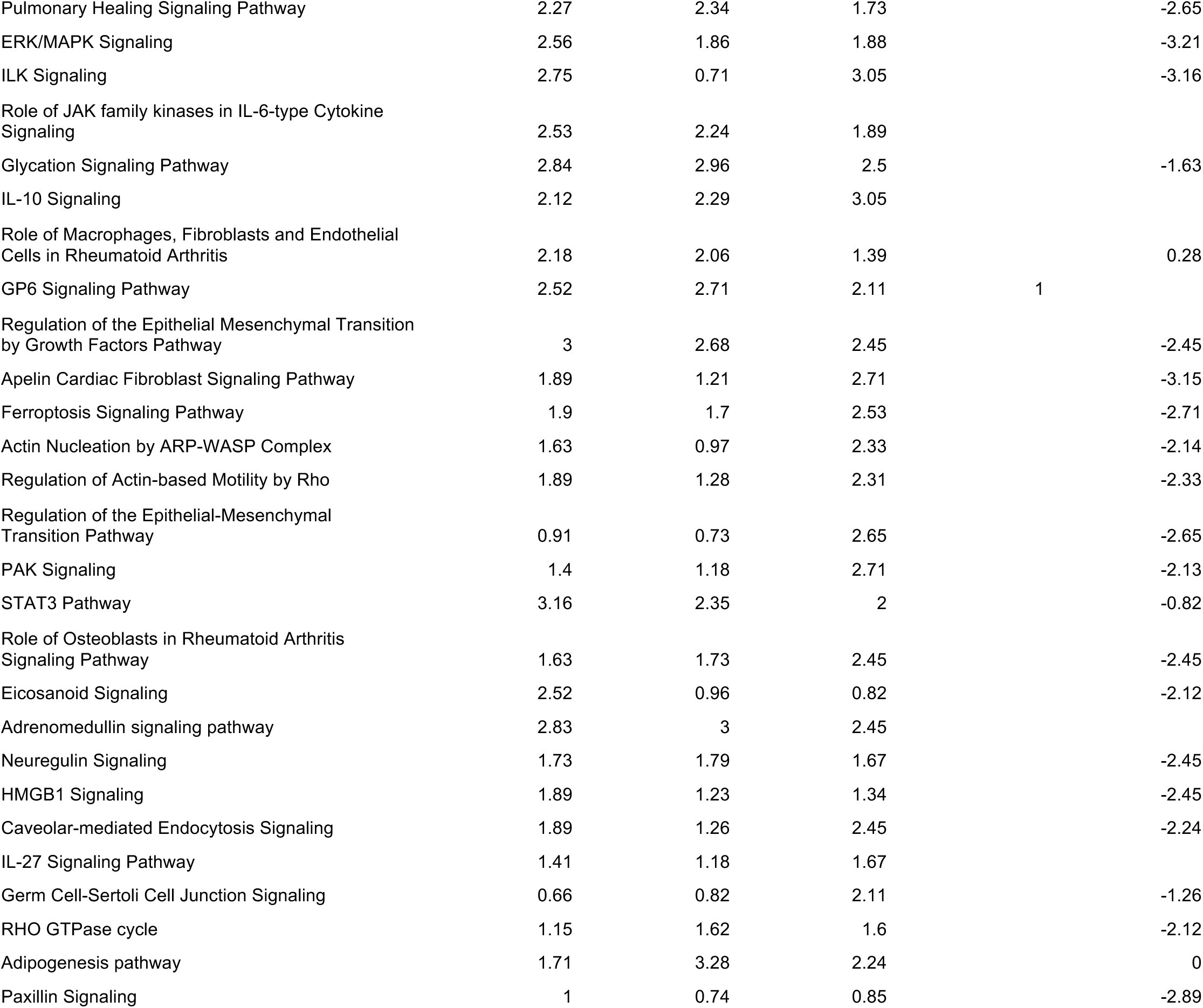
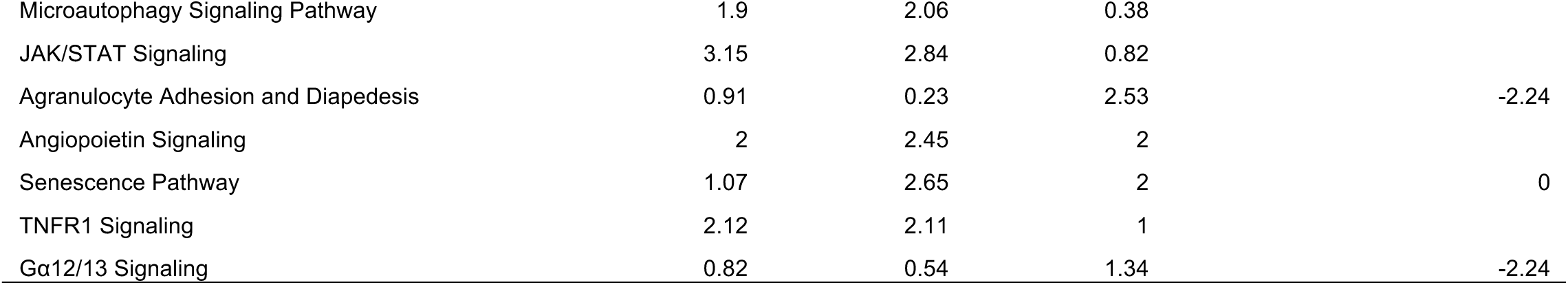
Activation z-scores across four gain-of-function disease states and the long-lived rescue for the conserved 59-pathway injury core.

### Supplementary Figures

**Figure S1.**
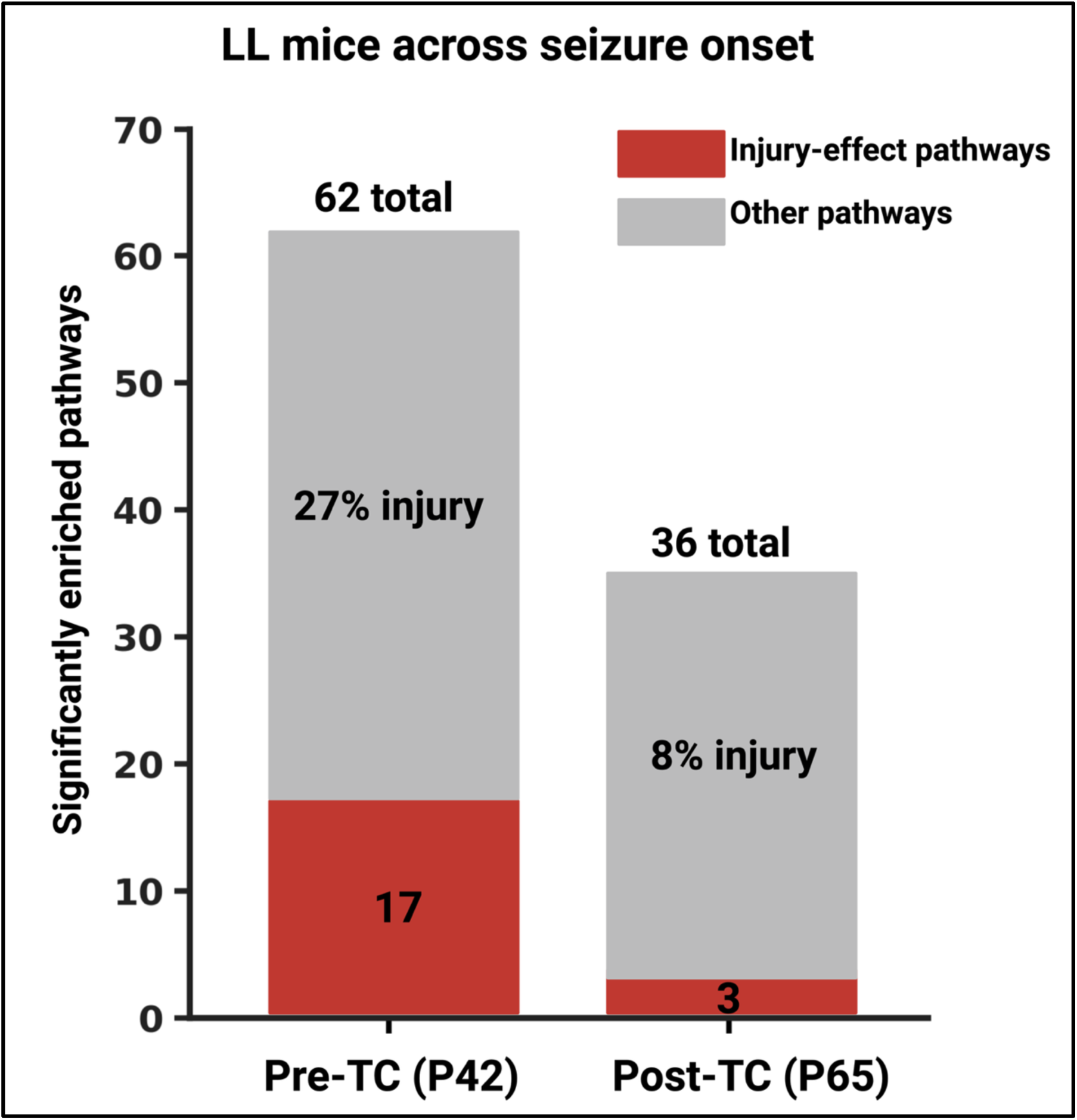
Durable suppression in the long-lived line. Significantly enriched pathways pre-TC (P42) and post-TC (P65); injury-effect pathways (red) decline from 27% to 8% of the enriched set after seizure onset. Enrichment presence only (activation z-scores unavailable for these contrasts). Created in BioRender. Bahramnejad, E. (2026) https://BioRender.com/27bg8yk

